# Atypical Cell Cycle Regulation Promotes Mammary Stem Cell Expansion and Therapeutic Resistance

**DOI:** 10.1101/2024.03.05.583524

**Authors:** Bre-Anne Fifield, John Vusich, Erika Haberfellner, Eran R. Andrechek, Lisa A. Porter

## Abstract

**Background:** The cell cycle of mammary stem cells must be tightly regulated to ensure normal homeostasis of the mammary gland to prevent abnormal proliferation and susceptibility to tumorigenesis. The atypical cell cycle regulator, Spy1 can override cell cycle checkpoints, including those activated by the tumour suppressor p53 which mediates mammary stem cell homeostasis. Spy1 has also been shown to promote expansion of select stem cell populations in other developmental systems. Spy1 protein is elevated during proliferative stages of mammary gland development, is found at higher levels in human breast cancers, and promotes susceptibility to mammary tumourigenesis when combined with loss of p53. We hypothesized that Spy1 cooperates with loss of p53 to increase susceptibility to tumour initiation due to changes in susceptible mammary stem cell populations during development and drives the formation of more aggressive stem like tumours.

**Methods:** Using a transgenic mouse model driving expression of Spy1 within the mammary gland, mammary development and stemness were assessed. These mice were intercrossed with p53 null mice to study the tumourigenic properties of Spy1 driven p53 null tumours, as well as global changes in signaling via RNA sequencing analysis.

**Results:** We show that elevated levels of Spy1 leads to expansion of mammary stem cells, even in the presence of p53, and an increase in mammary tumour formation. Spy1-driven tumours have an increased cancer stem cell population, decreased checkpoint signaling, and demonstrate an increase in therapy resistance. Loss of Spy1 decreases tumor onset and reduces the cancer stem cell population.

**Conclusions:** This data demonstrates the potential of Spy1 to expand mammary stem cell populations and contribute to the initiation and progression of aggressive, drug resistant breast cancers with increased cancer stem cell populations.

## Introduction

The mammalian mammary gland is a hormone-sensitive organ that undergoes complex structural and functional changes throughout life to permit lactogenesis. During puberty, pregnancy and through estrus, cell cycle activators support cell proliferation and expansion of the mammary epithelium [1–5]. Mammary cells are guided by morphological cues to form a branching ductal network with proliferation balanced by apoptosis as well as differentiation down luminal or myoepithelial lineages [6, 7]. A perfect balance of these events is required to ensure proper structure and organization of the mature, functional adult mammary gland. Aberrant regulation of mediators controlling these events may initiate or support tumourigenesis.

**<u>Ma</u>**mmary **<u>s</u>**tem **<u>c</u>**ells (MaSCs) and progenitor populations within the gland have been proposed as the cells of origin for many breast cancers [8–10]. Tight regulation over the expansion and differentiation of MaSCs throughout mammary development depends on the production and destruction of cell cycle mediators as well as tumour suppressors such as p53. Loss of p53 promotes a switch from asymmetric to symmetric division of stem and progenitor cells, and results in susceptibility to tumourigenesis [11–14]. Resolving the molecular mediators of mammary stem and progenitor populations throughout development is key to understanding how breast cancer initiates and progresses.

The atypical cell cycle mediator, Spy1 (also referred to as SpeedyA1, RINGO; gene *SPDYA*), is tightly regulated throughout mammary development, with protein levels being high during proliferative and regenerative phases and downregulated during differentiation [15]. Spy1 promotes cell cycle progression through both G1/S and G2/M by uniquely binding and activating both **<u>C</u>**yclin **<u>D</u>**ependent **<u>K</u>**inase (Cdk)1 and Cdk2 [16–18]. Unlike classical cyclin proteins, Spy1 activates Cdks independent of canonical post-translational modifications including phosphorylation of the Cdk T-loop by the Cyclin Activated Kinase (Cak) [19]. Previous work has elucidated a novel negative feedback loop in healthy cells between Spy1 and p53, where p53 promotes Spy1 degradation and supports a cell cycle arrest [20]. Sustained Spy1 expression occurs in the absence of p53 and overrides cell cycle checkpoints and promotes oncogenic transformation of human cell lines [21–25]. Driving Spy1 with a MMTV promoter on a mammary tumour resistant background (B6CBAF1/J) increases susceptibility to carcinogen-induced mammary tumorigenesis [20]. This occurs, at least in part, by overriding p53-mediated checkpoints allowing for accumulation of DNA damage, hence supporting that a Spy1-p53 feedback loop plays a critical role in mediating breast cancer susceptibility [20]. Spy1 protein levels are elevated in cases of human breast cancer as well as in other cancers including brain, liver, ovarian, and several blood cancers [21, 26–30] and elevated levels of Spy1 drives more aggressive, drug resistant disease in both **<u>E</u>**strogen **<u>R</u>**eceptor (ER) positive and Triple Negative Breast Cancer [26, 31]. Several publications have implicated Spy1 in expanding normal and cancer stem cell populations in the brain [28, 32, 33]. Recently, Spy1 has been shown to activate the epigenetic regulator EZH2 and support reprogramming to a pluripotent state [34]. Whether Spy1-mediated effects on stemness have any implications in the mammary gland was not previously addressed.

Herein we provide the first data demonstrating that overexpression of Spy1 within the mouse mammary gland alters mammary ductal development and promotes expansion of the MaSC population. We have previously shown that Spy1 cooperates with loss of p53 to increase susceptibility to tumourigenesis and confirm this finding on a mouse background sensitive to mammary tumourigenesis. Dissecting this further, we provide evidence that Spy1 driven tumours have increased stem cell and cell cycle progression signaling contributing to increased therapy resistance, linking early changes in mammary development to tumourigenic properties. This work adds to the growing literature supporting a role for this atypical cell cycle mediator in mammary gland development and breast tumorigenesis. More broadly this work provides a novel mechanistic connection between expansion of MaSC populations and breast cancer initiation, progression, and response to therapy.

## Materials and Methods

### Generation and Maintenance of MMTV-Spy1 Transgenic Mice

MMTV-Spy1 transgenic mice were generated as described [20] and backcrossed 10 generations onto an FVBN/J background. Routine genotyping was performed as described [20]. The p53 knockout mouse, B6.129S2-Trp53tm1Tyj/J, was purchased from Jackson Laboratory (002101) [35] and backcrossed 10 generations onto an FVBN/J background. Mice were maintained following the Canadian Council of Animal Care guidelines under animal utilization protocol 20-15 approved by the University of Windsor. The maximum allowed tumour volume of 1.5cm^3^ was not exceeded in any of the described studies according to protocol guidelines.

### Mammary Fat Pad Transplantation and Limiting Dilution Assay

For limiting dilution assays of normal mammary epithelial cells, mammary epithelial cells were isolated from 8-week-old and 6-month-old MMTV-Spy1 or FVBN/J littermate control mice as described [36]. Cells were transplanted into the cleared glands of 3-week-old FVBN/J mice and successful clearing was monitored via the addition of a cleared gland with no injected cells. For limiting dilution assays of primary tumour cells, cells were transplanted into the inguinal mammary glands of 6-week-old FVBN/J mice. Mammary epithelial cells from intercrossed MMTV-Spy1 and p53 null 8-week-old mice were transplanted into the cleared glands of 3-week-old FVBN/J mice for tumour studies and successful clearing was monitored via addition of cleared gland with no injection. Analysis was performed as described [37].

### Whole Mount Analysis

Inguinal mammary glands were placed on positively charged slides and incubated in Clarke’s Fluid (75% ethyl alcohol 25% acetic acid) overnight. The following day, slides were placed in 70% ethyl alcohol for 30 minutes and stained for a minimum of overnight in carmine alum (0.2% carmine, 0.5% potassium aluminium sulphate). Glands were placed in destain solution (1%HCl, 70% ethyl alcohol) for 4 to 6 hours and dehydrated in a series of ascending alcohol concentrations (70%, 95%, 100%) for 15 minutes each prior to clearing of the gland overnight in xylene. Slides were mounted with Permount Toluene (Fisher Scientific) solution prior to imaging.

### Histology and Immunostaining

Tissue was collected and fixed in 10% neutral buffered formalin and immunohistochemistry was performed as described [38]. Primary antibodies were as follows: Spy1 (1:200; PA5-29417; Thermo Fisher Scientific), PCNA (1:500; sc-9857; Santa Cruz); cleaved caspase-3 (1:250; 9661; Cell Signaling), cytokeratin 8 (1:100; MABT329 Millipore), phospho-histone H3 S10 (1:300; ab5176 Abcam) and smooth muscle actin (1:250; A2547 Sigma). Slides were imaged using the LEICA DMI6000 inverted microscope with LAS 3.6 software.

### Quantitative Real Time PCR Analysis

RNA was isolated using the Qiagen RNeasy Plus Kit following manufacturer’s instructions. Equal amounts of cDNA were synthesized using Quanta qScript Master Mix as per manufacturer’s instructions. Real time PCR was performed using SYBR Green detection (Applied Biosystems) and was performed and analysed using Viia7 Real Time PCR System (Life Technologies) and software.

### Primary Mammary Epithelial Cell Culture

Mammary epithelial cells (MECs) were isolated from inguinal mammary glands as previously described [36] and cultured at 5% CO2 at 37°C in DMEM/F12 supplemented with 10% FBS, 5µg/mL insulin, 1µg/mL hydrocortisone, 10ng/mL EGF and 1% P/S for adherent culture, and were sub-cultured using 0.05% trypsin. For mammosphere formation assays, cells were seeded at a density of 10,000-20,000 cells per mL of culture media (DMEM/F12 supplemented with 20ng/mL bFGF, 20ng/mL EGF, B27, 100µg/mL gentamicin, 1%P/S) in ultra low attachment plates. Mammospheres were dissociated with 0.05% trypsin to obtain single cell suspensions and seeded at a density of 1,000-5,000 cells/mL for secondary sphere formation. Transfection of primary MECs with sip53 (Santa Cruz) and siRNA control (Santa Cruz) was performed using siRNA Transfection Reagent as per manufacturer’s instructions. For treatment assays, cells were seeded at a density of 20,000 cells per well in a 24 well plate and treated with vehicle control (DMSO), 25nM doxorubicin, 6mM cyclophosphamide, 100nM paclitaxel, 10nM dinaciclib or 500nM palbociclib.

### PKH Staining

Primary MECs were stained with PKH26 dye (Sigma) as per manufacturer’s instructions and seeded for mammosphere formation assays to obtain primary mammospheres. Primary mammospheres were dissociated into single cell suspensions and sorted based on PKH26 staining intensity using a BD FACSARIA Fusion flow cytometer where single cell suspensions were then seeded for secondary sphere formation.

### RNA-Sequencing and Gene Expression Analysis

Total RNA was isolated using the Qiagen RNeasy Plus Kit following manufacturer’s instructions. RNA was sent to Psomagen and quality measured using Agilent 2100 Expert Eukaryote Total RNA NanoDrop. RNA libraries were prepared using TruSeq Stranded mRNA Library Prep Kit. Libraries were sequenced using the S4 chipset on the NovaSeq6000 instrument (Illumina) for 150bp paired end sequencing and 60M total reads. Raw FASTQ files were processed using the nf-core RNAseq pipeline (nf-core/rnaseq v3.12.0) [39, 40]. Adapter and quality trimming was performed using Trim Galore v0.6.7 [41]. FASTQ reads were aligned to the mouse GRCm39 reference genome using STAR [42]. RNA sequencing transcripts were quantified using Salmon v1.10.1 [43]. Quantified transcripts were analyzed using the nf-core differential abundance pipeline (nf-core/differentialabundance v1.4.0) [39, 40]. Differential expression analysis was performed using DESeq2 v1.34.0 and EnhancedVolcano v1.20.0 from Bioconductor in R. GSEA v4.3.2 was used for gene set enrichment analysis [44]. Gene expression plots were made using ggplot2 in R [45].

### Cell Culture

MCF7 (HTB-22; ATCC) were cultured in Dulbecco’s modified Eagle’s medium (DMEM; D5796; Sigma Aldrich) supplemented with 10% foetal bovine serum (FBS; F1051; Sigma Aldrich) and 1% P/S at 37°C and 5% CO2. MCF7 and primary MECs were lentivirally infected in serum and antibiotic free medium containing 8µg/mL polybrene. The pEIZ plasmid was a kind gift from Dr. B. Welm, and the pEIZ-Flag-Spy1 vector was generated as previously described [28]. The pLKO, pLKO-Spy1.1, pLKO-shSpy1.2, and pLB, pLB-shSpy1 plasmids were generated as described [16, 28]. The p53-GFP backbone was purchased from Addgene (11770) (p53-GFP was a gift from Geoff Wahl (Addgene plasmid #11770)), shp53 pLKO.1 puro was purchased from Addgene (19119) (shp53 pLKO.1 puro was a gift from Bob Weinberg (Addgene plasmid # 19119 ; http://n2t.net/addgene:19119 ; RRID:Addgene_19119)) [46].

### Statistics

A Mann-Whitney was performed for tumour studies. For all other data, a Student’s T-Test was performed. For experiments involving mouse samples and primary MECs, unequal variance was assumed. For cell line-based experiments, equal variance was assumed. For differential expression analysis, DESeq2 calculates p-values using the Wald test and BH-adjusted p-value (padj) is calculated using the Benjamini Hochberg method [47]. All experiments include at least 3 biological replicates and are representative of at least 3 experimental replicates.

## Results

### Spy1 increases ductal branching

The MMTV-Spy1 transgenic mouse model previously generated on a B6CBAF1/J background did not have any gross morphological differences in mammary development, despite a significant increase in mammary tumour susceptibility [20]. This was likely due to the nature of the model background as this strain is known to be more resistant to alterations in mammary development [48, 49]. To address this, the MMTV-Spy1 model was backcrossed 10 generations onto an FVBN/J background, a more commonly used strain for studying mammary development. Levels of Spy1 mRNA and protein were confirmed to be significantly upregulated in MMTV-Spy1 mice as compared to littermate controls (Figure S1 A,B). Previous data has demonstrated that elevation of Spy1 in HC11 cells was able to promote accelerated ductal development upon mammary fat pad transplantation [15]. Rate of ductal elongation and number of branch points was quantified throughout pubertal development. MMTV-Spy1 mice had a significant increase in ductal elongation at 8 weeks (Figure S1C) and a significant increase in number of branch points across all time points collected (Figure S1D: upper graph). A significant increase in the overall percent fat pad filling at all time points was also observed (Figure S1D: lower graph). These alterations in ductal branching corresponded to a significant increase in the percentage of PCNA positive cells at all time points examined, and a significant decrease in cleaved caspase 3 (CC3) at 8 weeks of age (Figure S2A,B). Thus, Spy1 significantly increases rates of proliferation and decreases apoptosis at select time points in the developing mouse mammary gland. This may be a contributing factor to the increased ductal branching noted.

Increased rates of proliferation may impact the cellular composition of the mammary gland by altering cell cycle dynamics leading to changes in the differentiation status of the cells. First, inguinal mammary glands from 8-week-old mice were assessed for luminal and myoepithelial markers **<u>c</u>**yto**<u>k</u>**eratin **<u>8</u>** (CK8) and **<u>s</u>**mooth **<u>m</u>**uscle **<u>a</u>**ctin (SMA) respectively. No change in expression of either marker was noted in the MMTV-Spy1 inguinal mammary glands (Figure S2C). Hormone receptor status was assessed via immunohistochemistry from 8 and 12-week-old MMTV-Spy1 mice and littermate controls. MMTV-Spy1 mice showed accelerated expression of **<u>p</u>**rogesterone **r**eceptor (PR) and ERα positive cells at 8 weeks of age (Figure S3). This collectively supports that there is an early expansion of the hormone receptor positive population in mammary glands with Spy1 elevation, and potential alterations in the differentiation status of the mammary gland.

### Elevated levels of Spy1 lead to expansion of the mammary stem cell population

An increase in proliferation coupled with alterations to the hormone receptor status in the mammary gland may impact the MaSC population, as this population is hormone receptor negative [50, 51]. To determine if Spy1 can promote expansion of the MaSC population, a limiting dilution assay was performed using **<u>m</u>**ammary **<u>e</u>**pithelial **<u>c</u>**ells (MECs) isolated from 8-week-old mice. Strikingly, the MMTV-Spy1 mice had more than a 10-fold increase in the stem cell population (Figure 1A). This increase was maintained over time as demonstrated using limiting dilution assays on MECs isolated from 6-month-old mice (Figure 1B). As a secondary measurement of the stem cell population, primary MECs were isolated from 10-12-week-old MMTV-Spy1 mice and their control littermates and a mammosphere formation assay was performed. MECs isolated from MMTV-Spy1 mice had a significantly higher **<u>s</u>**phere **f**ormation **<u>e</u>**fficiency (SFE) as compared to control MECs when assessing the formation of primary spheres (Figure 1C). Furthermore, a higher rate of SFE was sustained when sequentially passaged, whereas mammospheres from littermate controls demonstrated a decline in SFE over time (Figure 1C). To further address the ability of Spy1 to promote stem cell characteristics in the mammary gland, primary MECs from MMTV-Spy1 and littermate control mice were isolated, stained with the label retaining dye PKH26 and seeded for a mammosphere formation assay. PKH26 is a fluorescent dye which binds cell membranes and segregates into daughter cells such that it can be used to identify proliferative history. It has been previously shown that mammary cells which retain high PKH26 staining contain a higher proportion of MaSCs as compared to PKH26 negative cells [11, 13, 52]. Following mammosphere formation, cells were sorted based on PKH26 staining and seeded as secondary mammospheres as PKH26^high^ and PKH26^negative^ populations (Figure 1D). As expected, PKH26^negative^ populations had a reduced SFE compared to PKH26^high^ populations, however MMTV-Spy1 mammospheres increased SFE significantly in both PKH26^negative^ and PKH26^high^ populations. Taken together, this suggests that elevated levels of Spy1 promote stem cell potential of MECs, leading to expansion of the MaSC population.

**Figure 1:**
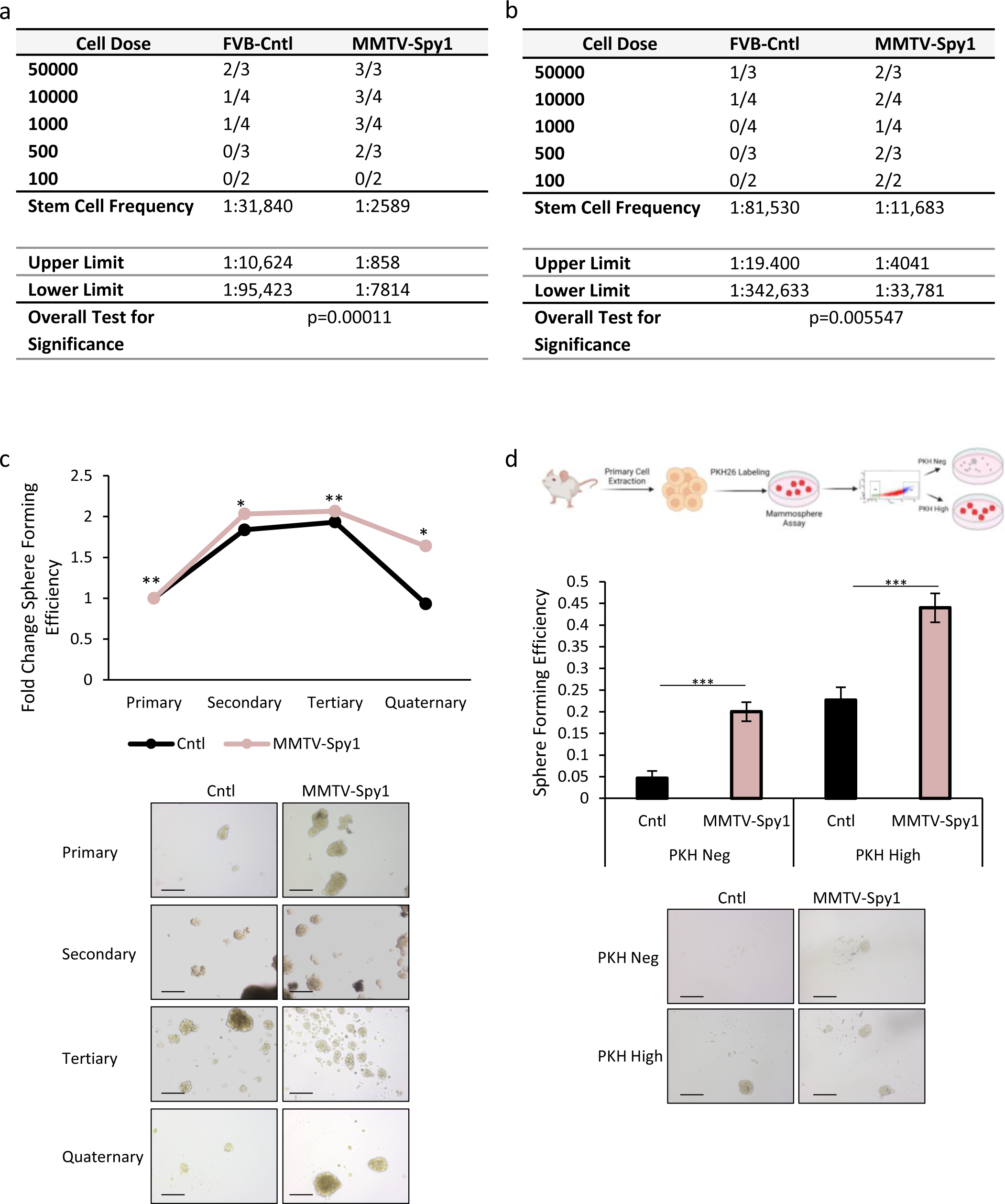
MMTV-Spy1 mice have an increased MaSC population. Limiting dilution assays were performed using primary cells from **a**) 8-week-old and **b**) 6-month-old littermate control (FVB-Cntl) and MMTV-Spy1 mice. **c**) Mammospheres from MMTV-Spy1 and littermate control mammary primary cells were serially passaged. Fold change in sphere forming efficiency is depicted (top panel; n=3). Representative images depicted in bottom panel. **d**) Primary cells were isolated from control and MMTV-Spy1 inguinal mammary glands, stained with PKH and cultured and sorted as depicted in schema in top panel. Sphere forming efficiency is graphically represented in middle panel. Representative images depicted in bottom panel. (n=6); Scale bar = 200 μm; Errors bars represent SE; Student’s T-test. *p<0.05, **p<0.01, ***p<0.001.

### Spy1 overrides p53 to promote expansion of the MaSC population

Spy1 can cooperate with the loss of p53 to promote susceptibility to mammary tumourigenesis, and functionally overrides p53 in response to various types of DNA damage [20, 24, 25]. Loss of p53 on its own leads to an increase in symmetric division and expansion of MaSC populations [11–14]. On the other hand, elevation of p53 supports a reduction in the MaSC population, asymmetric division and an increase in differentiation [11–13]. To determine if Spy1 plays a role in p53-mediated effects in the MaSC population, primary MECS from MMTV-Spy1 and littermate control mice were cultured as mammospheres in the presence and absence of Nutlin3, a small molecule that inhibits MDM2 dependent degradation of p53 leading to p53 stabilization [53]. SFE in control mammospheres was significantly reduced in the presence of Nutlin3 whereas no effect was seen on MMTV-Spy1 mammospheres (Figure 2A), thus indicating that Spy1 can override p53 to allow for continued expansion of the MaSC population. This effect was also seen in MCF7 cells manipulated with lentivirus to overexpress Spy1 (Figure 2B).

**Figure 2:**
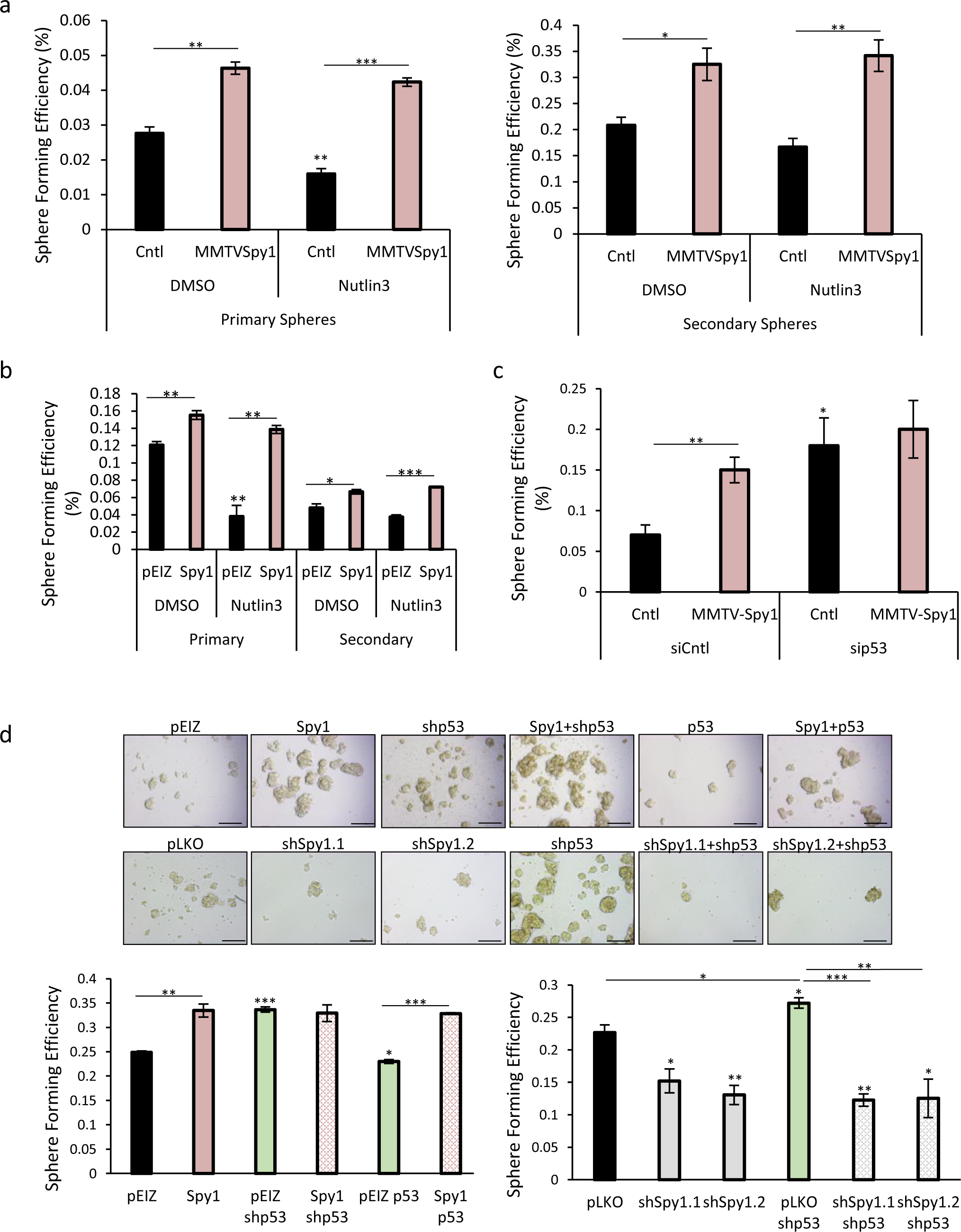
Spy1 overrides p53 to promote MaSC expansion. **a**) Primary mammary epithelial cells were isolated from inguinal mammary glands of MMTV-Spy1 and littermate control mice and cultured as mammospheres in the presence and absence of 2.5µM Nutlin3. Sphere forming efficiency of primary mammospheres (left panel) and secondary mammospheres (right panel) is depicted (n=3). **b**) Control (pEIZ) or Spy1 overexpression (Spy1) MCF7 cells were treated with 2.5µM Nutlin3 and cultured as mammospheres. Sphere forming efficiency in the presence or absence of Nutlin3 is depicted. **c**) siRNA mediated knockdown of p53 was performed in primary mammary epithelial cells from control and MMTV-Spy1 inguinal mammary glands, and cells were cultured as mammospheres. Sphere forming efficiency is depicted (n=5). **d**) MCF7 cells were lentivirally infected with pEIZ, pEIZ Spy1 (Spy1), pLKO, pLKO shSpy1 (shSpy1.1, shSpy1.2), p53 or shp53. MCF7 cells with Spy1 overexpression (left panel) or Spy1 knockdown (right panels) in the presence or absence of p53, were cultured as mammospheres. Sphere forming efficiency was assessed and quantified (lower panels). Representative images shown in upper panels. Scale bar = 200 μm; n=3; Errors bars represent SE; Student’s T-test. *p<0.05, **p<0.01, ***p<0.001.

To assess the effects of loss of p53 in Spy1 overexpressing samples, primary MECs were again isolated from MMTV-Spy1 and littermate control mice and p53 was knocked down via siRNA and cells were seeded for mammosphere formation. As expected, knockdown of p53 increases SFE, however, while MMTV-Spy1 alone increased SFE, the combination of MMTV-Spy1 with sip53 did not result in a significant increase in SFE over that of sip53 alone (Fig 2C). This suggests that Spy1 effects on MaSC expansion are via the ability to override p53. To further validate this result, MCF7 cells were manipulated with lentivirus to overexpress and knockdown both Spy1 and p53 in varying combinations. Spy1 was able to promote SFE in the presence of elevated p53 (Figure 2D) and was not able to significantly increase SFE with loss of p53 (shp53) over that of shp53 alone (Figure 2D). Interestingly, knockdown of Spy1 alone lead to a significant decrease in SFE and when combined with knockdown of p53, completely ablated the effects of loss of p53 on SFE indicating that in MaSCs with loss of p53, the ability to expand this population may be dependent on Spy1 (Figure 2D). Taken together this data demonstrates that Spy1 can override p53 to promote MaSC expansion and may be required for promoting MaSC division with loss of p53.

### Loss of p53 does not alter Spy1 mediated effects on normal mammary development

Despite alterations to ductal branching, early increases in hormone receptor positive cells and an increase in the MaSC population, only 2 spontaneous mammary tumours were documented over 50 MMTV-Spy1 mice allowed to age up to 2 years of age. Previous data has shown that Spy1 cooperates with loss of p53 to promote mammary tumourigenesis; however, whether the changes observed with Spy1 elevation during development, particularly with the MaSC population, contribute to this has not yet been explored. To address this, MMTV-Spy1 mice were intercrossed with p53 null mice. First, inguinal mammary glands were collected at 8 weeks to determine if the loss of p53 altered any of the developmental phenotypes noted in the MMTV-Spy1 model. Loss of p53 did not affect the ability of Spy1 to promote ductal elongation or increase the number of branch points (Figure S4), and no changes in differentiation status were observed with CK8 and SMA (data not shown). Previous work has characterized a positive feedback loop between ERα and p53 [54, 55]. Additionally, there was no change on the ability of Spy1 to increase PR or ERα expression at 8 weeks in the absence of p53 (Figure S5). Collectively, these data support that Spy1 effects on mammary development are independent of p53 status.

### Spy1 p53 null tumours have elevated stem cell and cell cycle signaling

While loss of p53 did not alter ductal development or differentiation status of the mammary gland in combination with elevated Spy1, Spy1 and p53 do appear to have a link with respect to the MaSC population. To address the impact of an increased MaSC population on Spy1 p53 null driven tumours, MMTV-Spy1 mice were again intercrossed with p53 null mice. Primary MECs were extracted at 8 weeks of age and mammary fat pad transplantation was performed in cleared glands of 3-week-old FVBN/J mice as described [20]. Similar to previous findings, Spy1 cooperated with partial or complete loss of p53 to increase HANs and tumour formation (Figure 3A). No significant difference in rate of tumour onset was found (Figure S6A). Immunohistochemical analysis revealed a significant increase the percentage of phospho-histone H3 positive cells and a significant decrease in CC3 positive cells in Spy1 p53 null tumours, indicating increased proliferation and decreased apoptosis as compared to p53 null tumours (Figure 3B-C).

**Figure 3:**
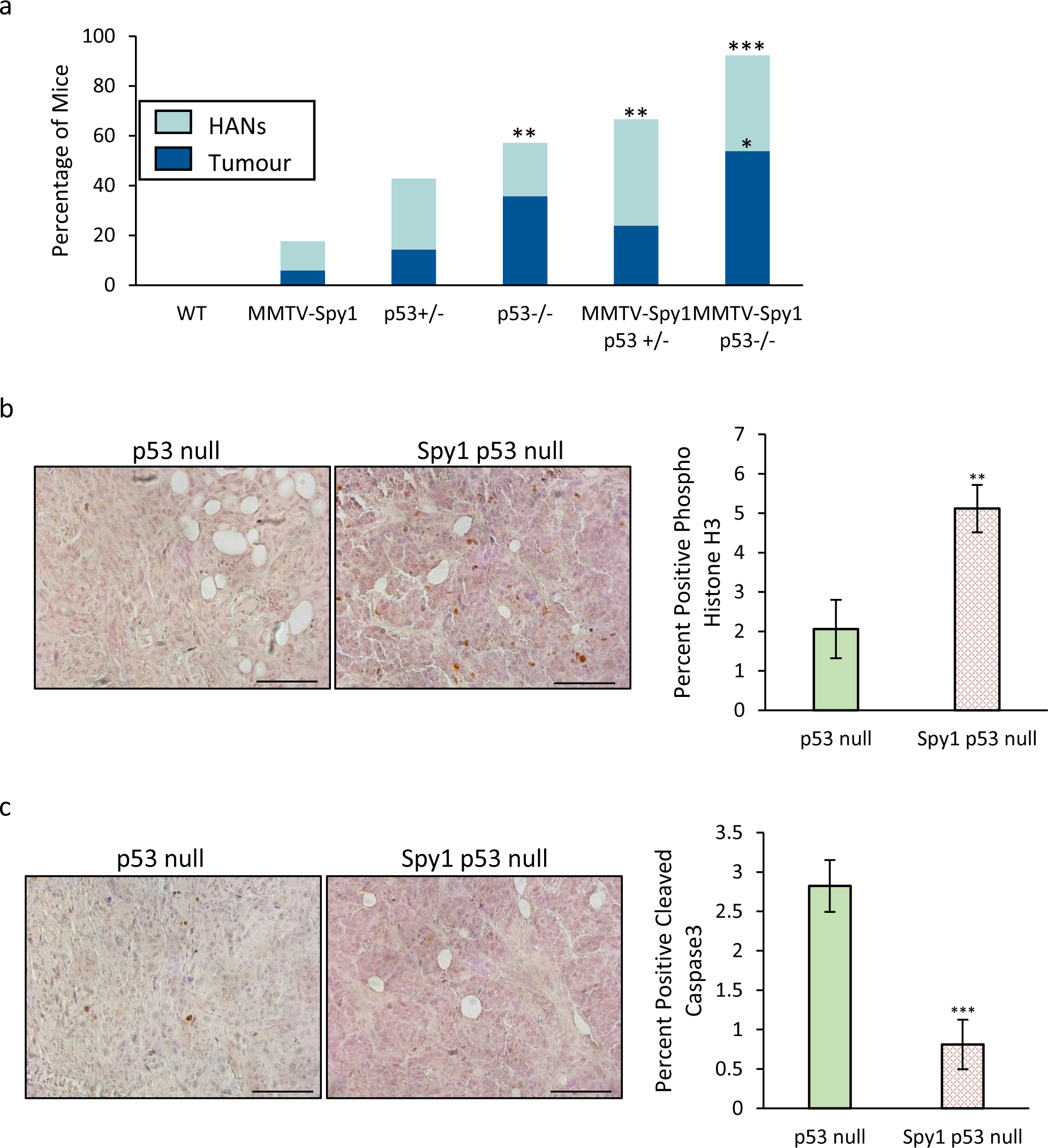
Elevated Spy1 increases mammary tumour formation. **a**) Primary mammary epithelial cells from MMTV-Spy1 and p53 null intercrossed mice were injected into the cleared fat pads of wildtype mice and monitored weekly for HANs and tumour development. HANs were only quantified in tumour negative mice (Cntl n=15, MMTV-Spy1 n=17, p53+/- n=21, p53-/- n=14, MMTV-Spy1 p53+/- n=21, MMTV-Spy1 p53- /- n=13). Immunohistochemical analysis of **b**) phospho histone H3 and **c**) cleaved caspase 3 in Spy1 p53 null and p53 null tumours (p53 null n=3; Spy1 p53 null n=4). Representative images shown in left panels where blue represents hematoxylin nuclear stain and brown represents **b**) phospho histone H3 or **c**) cleaved caspase 3. Quantification of percent positive **b**) phospho histone H3 or **c**) cleaved caspase 3 depicted graphically in right panels. Scale bar = 100 μm; Mann-Whitney (a) and Student’s T-test (b&c). *p<0.05, **p<0.01, ***p<0.001.

Histological analysis revealed that the Spy1, p53 null and Spy1 p53 null tumours shared close histological features within groups rather than shared features between groups (Figure S6B), suggesting differences in tumour subtypes and cellular composition. Differences in histology may point to alterations in key signaling pathways between p53 null and Spy1 p53 null driven tumours. To further elucidate these differences, RNA sequencing (RNAseq) followed by **<u>G</u>**ene **<u>S</u>**et **<u>E</u>**nrichment **<u>A</u>**nalysis (GSEA) [44] was performed on p53 null and Spy1 p53 null tumours. GSEA revealed upregulation of pathways associated with branching in mammary gland duct morphogenesis, as well as elevated breast cancer and mammary luminal progenitor pathways (Figure 4, S7, S8). Pathways associated with breast cancer progression and increased cell cycle signaling were also upregulated (Figure 4, S7-S9). Thus, this analysis revealed commonalities in upregulated signaling pathways associated with changes observed in the normal mammary gland that may be contributing factors to susceptibility to tumourigenesis.

**Figure 4:**
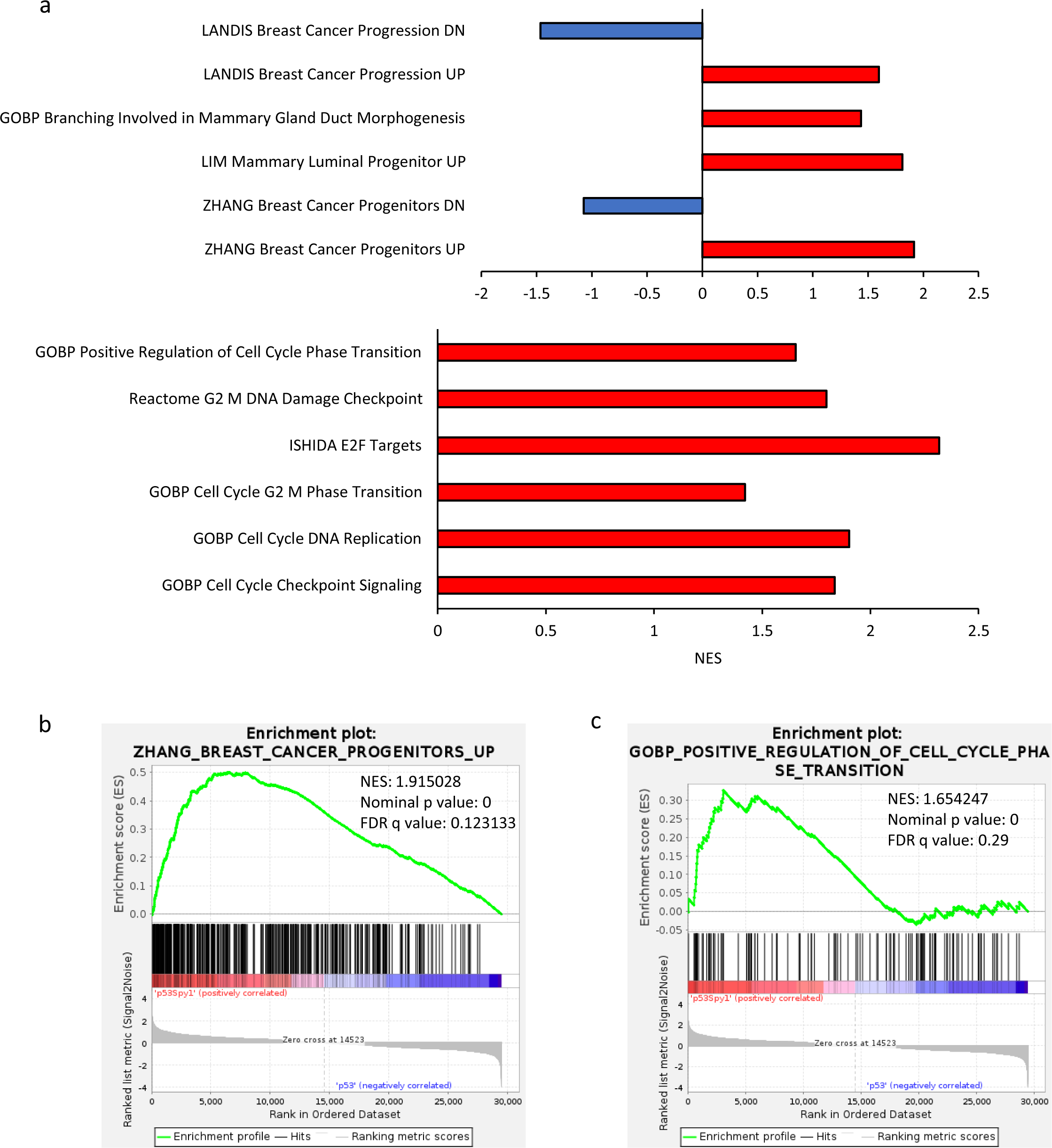
Spy1 driven tumours have unique signaling pattern. GSEA was performed on Spy1 p53 null versus p53 null driven tumours. **a**) Normalized enrichment scores (NES) of selected gene sets are depicted graphically. **b**) Enrichment plots of breast cancer progenitor gene set (left panel) and positive regulation of cell cycle phase transition gene set (right panel) are depicted.

When assessing changes in gene expression, approximately 500 genes were significantly upregulated while over 600 genes were significantly downregulated in Spy1 p53 null tumours as compared to p53 null alone (Figure 5A). Among the top genes significantly upregulated was Chi3l1 (Chil1) (Figure 5B), a gene associated with poor prognosis in breast cancer [56]. In addition, other genes associated with stem cell populations or changes seen in MMTV-Spy1 mice during normal mammary development were also impacted. FoxA1, a known driver of stem cell proliferation and regulator of ERα expression [57], as well Prom1, a regulator of luminal cell differentiation and ductal branching [58] were also significantly upregulated (Figure 5C,D). Significantly downregulated was Cdkn1a, a target of p53 and inhibitor of MaSC expansion (Figure 5E). To validate these findings, expression of Chi3l1, FoxA1, Prom1 and Cdkn1A were examined via qRT-PCR in Spy1 p53 null and p53 null tumours as well as MMTV-Spy1 p53 null intercrossed inguinal mammary glands from 8-week-old mice (Figure S10). qRT-PCR analysis of tumour samples confirmed the findings from RNAseq, and similar trends were seen in p53 null and MMTV-Spy1 p53 null 8-week-old inguinal mammary glands (Figure S10), thus demonstrating that early changes in normal development may contribute to the onset of tumourigenesis and tumour characteristics.

**Figure 5:**
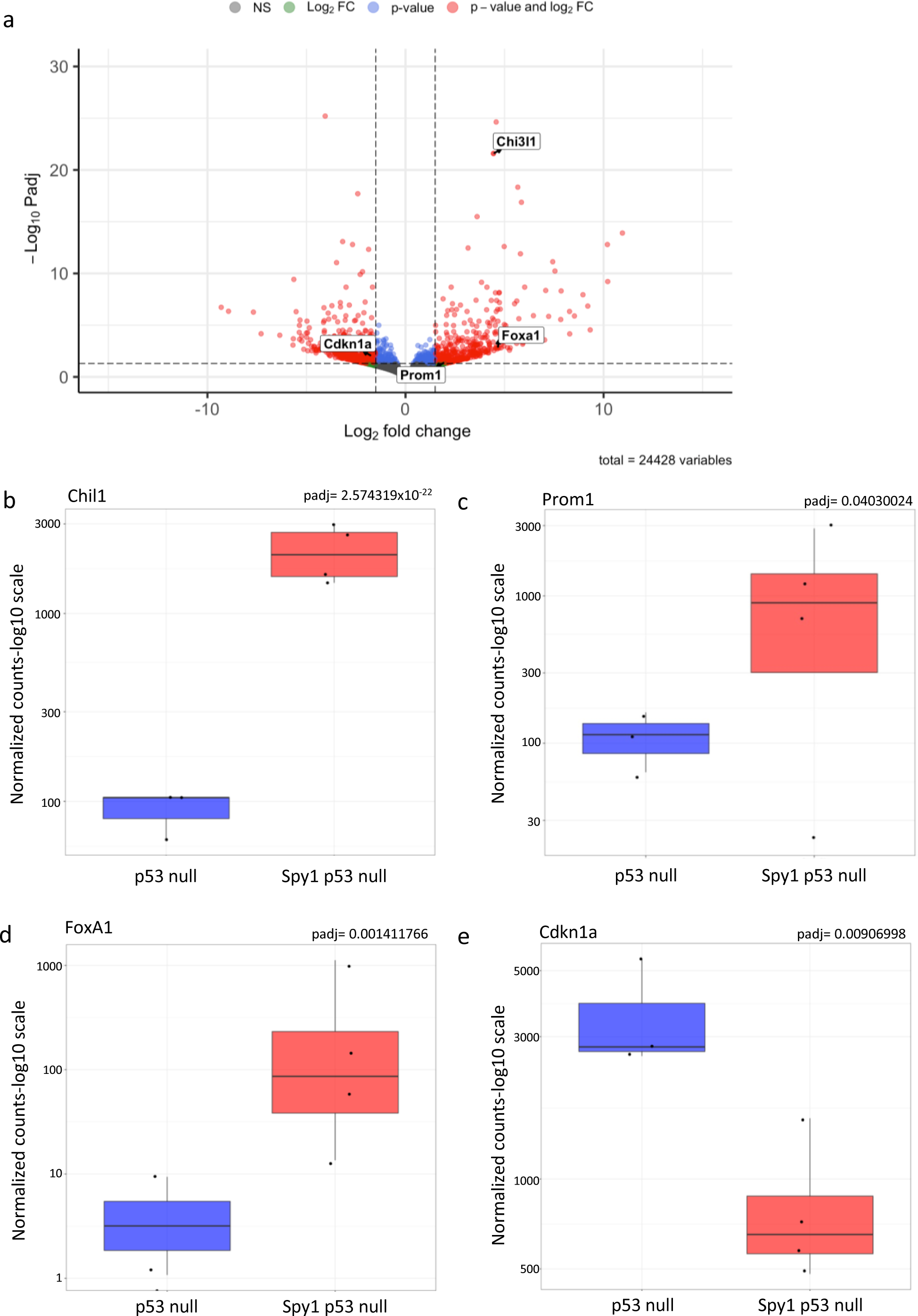
Changes in gene expression reflect gene set and developmental changes with elevated Spy1 expression. **a**) Volcano plot depicting differentially expressed genes in Spy1 p53 null as compared to p53 null tumours. Normalized gene expression counts of **b**) Chil1 (Chi3l1), **c**) Prom1, **d**) FoxA1 and **e**) Cdkn1a from RNA sequencing data with BH-adjusted p-value from DESeq2 indicated in the top right of each graph.

### Expansion of the breast cancer stem cell population with loss of p53 and elevated Spy1 drives therapeutic resistance

Given the changes in signaling pathways associated with stem cell populations seen in Spy1 p53 null tumours, the stem cell population was assessed to determine if tumours arising in glands with elevated Spy1 had an increased proportion of stem cells. Primary tumour cells were isolated and cultured in non-adherent conditions to test SFE. When comparing cells from p53 null and MMTV-Spy1 p53 null tumours, a significantly higher SFE was found in the MMTV-Spy1 p53 null tumours as compared to p53 null tumours alone (Figure 6A). A higher rate of SFE was seen when examining secondary sphere formation, although this effect was not significant (Figure 6A). In addition, mammospheres from Spy1 tumours were significantly larger than mammospheres from any tumours with p53 alteration (Figure S11). A limiting dilution assay was performed on primary tumour cells from Spy1, p53 null and Spy1 p53 null tumours as a gold standard test to the observed increase in the **<u>b</u>**reast **<u>c</u>**ancer **<u>s</u>**tem **<u>c</u>**ell (BCSC) population. p53 null tumours had the lowest population of BCSCs as compared to Spy1 alone and Spy1 p53 null tumours (Figure 6B). Taken together, this data demonstrates that Spy1 leads to expansion of not only the normal MaSC population, but also the BCSC population.

**Figure 6:**
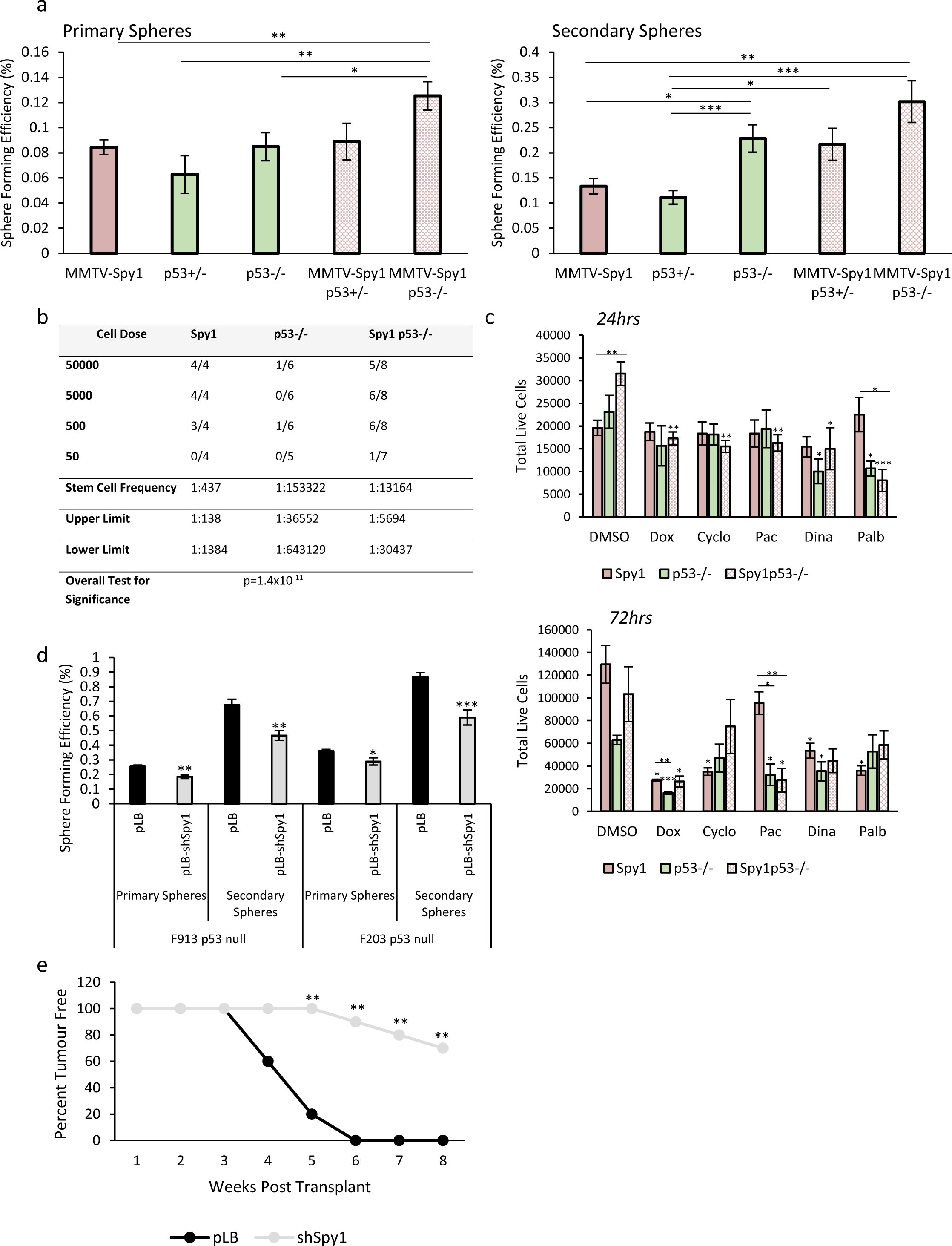
Loss of Spy1 delays tumour onset. **a**) Cells were isolated from tumours developed following mammary fat pad transplantation and mammosphere formation assays were performed. Sphere forming efficiency was quantified in primary (left panel) and secondary (right panel) mammospheres (n=3). **b**) Primary cells from Spy1, p53 null and Spy1 p53 null tumours were injected into fat pads of wildtype mice for a limiting dilution assay. **c**) Primary tumour cells from Spy1, p53 null and Spy1 p53 null tumours were treated with a panel of therapeutic agents for 24 hours (top panel) and 72 hours (bottom panel). Trypan blue exclusion analysis was performed to assess total number of live cells (n=3). **d**) Primary cells from p53 null tumours were infected with lentivirus containing control (pLB) or Spy1 knockdown (pLB-shSpy1) and cultured as mammospheres. Sphere forming efficiency was assessed and quantified for primary and secondary spheres (n=3). **e**) p53 null primary tumour cells infected with control (pLB) or knockdown of Spy1 (pLB-Spy1) were injected into fat pad of FVBN/J mice. Timing of tumour onset is depicted (n=10). Errors bars represent SE; Student’s T-test (a, c), Mann-Whitney (e). *p<0.05, **p<0.01, ***p<0.001.

Spy1 driven tumours have a significant increase in the BCSC population, and increased cell cycle progression signaling. The BCSC population is hypothesized to be a driving force in driving breast tumour growth and relapse [10], due in part to drug resistance, and this combined with alterations in cell cycle progression, may point to a role for Spy1 in driving drug resistance. To test this, primary tumour cells from Spy1, p53 null and Spy1 p53 null tumours were treated with a variety of chemotherapeutic agents and inhibitors that target various aspects of the cell cycle machinery for 24 and 72 hours. While the trends for each agent used varies, in all cases those tumours with elevated Spy1 displayed the largest increase in drug resistance over p53 null tumours alone (Figure 6C).

### Loss of Spy1 delays tumour formation and decreases the BCSC population

To determine if Spy1 is essential for expanding the BCSC population in p53 null tumours, Spy1 was knocked down using lentiviral manipulation in two p53 null primary tumour cell lines. Manipulated cells were then seeded for mammosphere formation assays. A decrease in Spy1 levels resulted in a significant decrease in SFE in both primary tumour cell lines in both primary and secondary mammospheres (Figure 6D). Next, control and Spy1 knockdown p53 null primary tumour cells were injected into inguinal mammary glands of 6-week-old mice and followed for tumour formation. The knockdown of Spy1 lead to a significant decrease in tumour formation as compared to control (Figure 6E). Thus, Spy1 cooperates with loss of p53 to drive the initiation and progression of drug resistant breast tumours with an increased BCSC population, which may be a contributing factor to therapy resistance and relapse (Figure 7).

**Figure 7:**
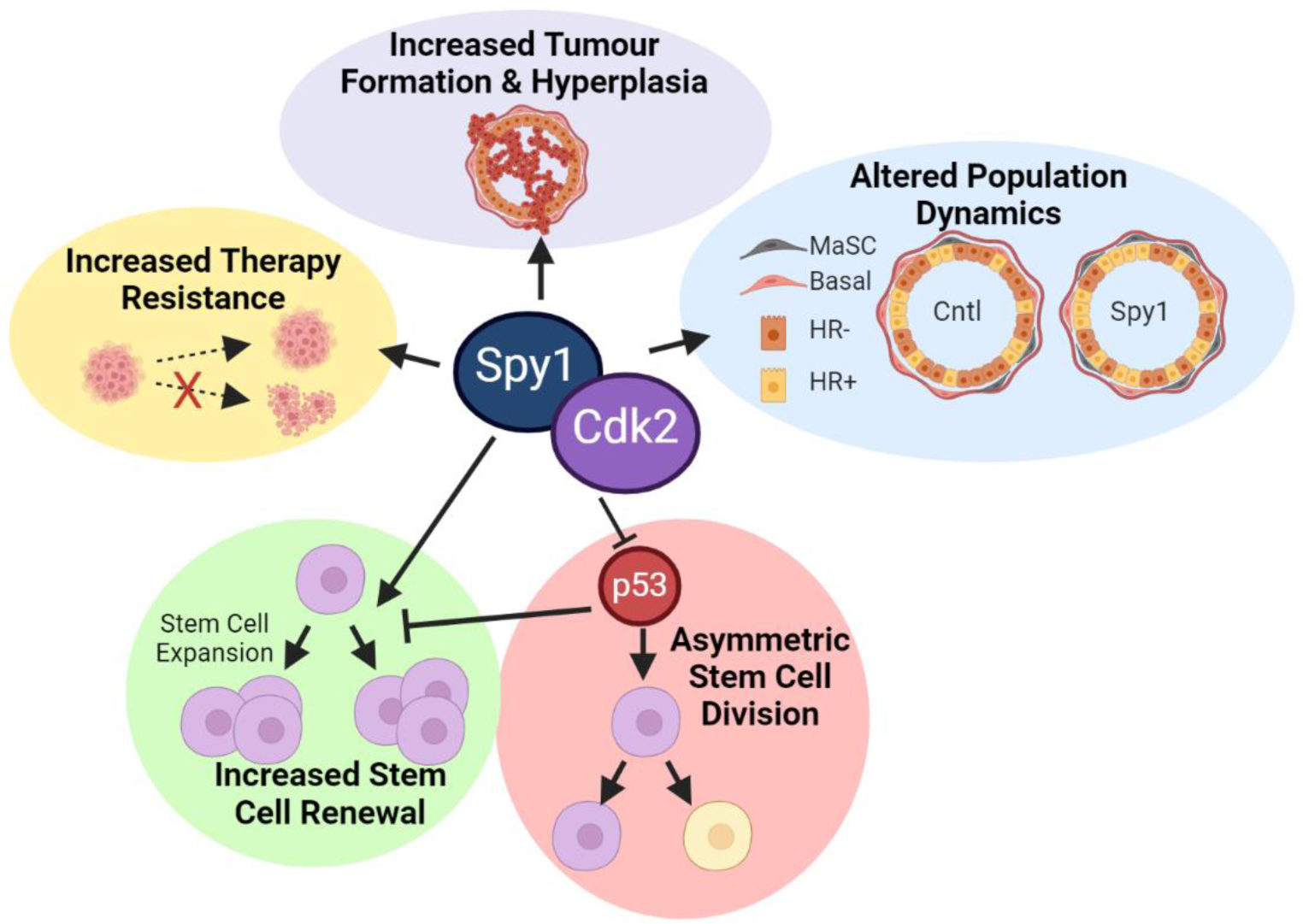
Summary of effects of elevated Spy1 on normal and abnormal development of the breast. Elevated Spy1 drives expansion of the MaSC population, in part by overriding p53, leading to initiation of tumours with an increase in the BCSC population. Spy1 p53 null tumours have a unique signaling signature over that of p53 null tumours which in part may contribute to increased therapeutic resistance.

## Discussion

Disruptions during normal mammary development that alter the ductal network and cellular composition of the mammary gland can have long lasting implications including increased susceptibility to mammary tumourigenesis. The transgenic MMTV-Spy1 mouse model revealed that sustained elevation of Spy1 can promote susceptibility to breast tumourigenesis by altering normal ductal development and increasing the MaSC population, a vulnerable population of cells thought to be the cell of origin for many types of breast cancer [8, 9]. Overexpression of Spy1 within the mouse mammary gland leads to increased ductal branching and enhanced ductal elongation, in part due to increased proliferation. While common luminal and basal differentiation markers remain unaltered, Spy1 accelerated the expression of hormone receptors. Previous work has shown that Spy1 mediated activation of ERK signaling leads to a significant increase in ligand independent phosphorylation and activation of ERα [31]. Whether the increase in ERα seen *in vivo* is mediated through a similar mechanism is an important next step of investigation. Given the important role hormone receptor signaling plays in regulating MEC proliferation, alterations in hormone receptor status may have profound implications on proliferative status and mutational accumulation over the lifetime of the mammary gland. Thus, the alteration in hormone receptor status and proliferative state of the mammary glands of MMTV-Spy1 mice may create an environment primed for oncogenesis.

In addition to alterations in hormone receptor status, the MaSC population was significantly increased in MMTV-Spy1 mice. Elevated levels of Spy1 were able to impart stemness properties to progenitor cell populations and increased the frequency of the MaSC population. MaSC populations are considered vulnerable cell populations to damaging events and may be the cell of origin for mammary tumourigenesis [8, 9]. Tumour suppressors such as p53 have tight control over expansion of this population, with p53 promoting asymmetric division, preventing dangerous expansion of this population [11]; however, Spy1 was able to override p53 and promote continued expansion of the MaSC population even in the presence of elevated p53. In addition, knockdown of p53 demonstrated that the enhanced stem cell expansion seen with loss of p53 requires Spy1 expression as the combined loss of Spy1 and p53 prevented continued expansion of the stem cell population. Further work is required to determine the exact mechanism by which this occurs and if Spy1 is expanding the MaSC population via alterations in asymmetric versus symmetric division, as has been seen in the brain tumour initiating cell population [28]. Further demonstrating the implications of an expanded MaSC population, the combined loss of Spy1 and p53 led to development of mammary tumours with an increased BCSC population and increased signaling associated with breast cancer progenitor populations. This demonstrates the critical role that maintenance of checkpoint signaling plays in restraining the MaSC population to prevent the expansion of a vulnerable cell population with potentially deleterious mutations that can act as the cell of origin in mammary tumourigensis. Tumours with elevated Spy1 had a significant increase in various mediators of breast cancer prognosis and normal mammary gland development that were also altered in samples obtained from MMTV-Spy1 p53 null intercrossed mice during puberty. Chi3l1 is associated with poor prognosis and has recently been shown to have a role in promoting an immunosuppressive tumour immune microenvironment in a mouse model of breast cancer [59]. Whether Spy1 may also be capable of contributing to an immunosuppressive state in mammary tumours is an important area of future study. Spy1 p53 null tumours also had a significant increase in FoxA1 and Prom1, both important mediators of normal mammary development and both implicated in BCSC populations [57, 58, 60, 61]. These changes in expression are linked to changes observed in mammary gland development of MMTV-Spy1 mice, as increased ductal branching and accelerated hormone receptor expression were observed and would be consistent with elevated FoxA1 and Prom1 expression [57, 58]. Importantly, Spy1 has previously been shown to expand Prom1 cancer stem cell populations in brain tumours [28] Given the evidence to support the existence of a Prom1 positive BCSC population [61], exploring whether Spy1 is selectively mediating this population through similar changes in symmetric division seen in Prom1 positive brain tumour initiating cell populations is important future work [28].

Analysis of the tumours revealed an increase in cell cycle signaling in tumours with elevated Spy1 and loss of p53 as compared to loss of p53 alone. Spy1 p53 null tumours were more resistant to various forms of therapy and had increased proliferative and decreased apoptotic capacity. This increase in tumour cell cycle signaling highlights the important role intact cellular checkpoints play in maintaining the integrity of the cell, and points to a potential role for Spy1 in the early stages of the initiation of tumourigenesis via disruption of cellular checkpoints and altered proliferative status. Restoration of cellular checkpoints to inhibit cellular proliferation and promote apoptosis has long been a goal of therapeutic intervention. This data supports that Spy1 could be a novel therapeutic target for a select subset of breast cancer patients with elevated levels of Spy1.

## Conclusions

Collectively, this data demonstrates that the atypical cell cycle regulator Spy1 plays an important role during normal mammary development. Overexpression of Spy1 alters the proliferative status of the mammary gland leading to acceleration of ductal elongation and increased ductal branching. Spy1 also alters the cellular composition of the mammary gland leading to altered hormone receptor status expression and expansion of the MaSC population. This data shows for the first time that the ability of Spy1 to override p53 can lead to changes in distinct cell populations with Spy1 able to override p53 driven asymmetric division of the MaSC population promoting increased expansion. This leads to increased susceptibility to mammary tumour formation with alterations in key checkpoint signaling pathways elucidating an important potential new therapeutic target for the treatment of select subsets of breast cancer patients.

## List of Abbreviations

BCSC: breast cancer stem cell
CC3: cleaved caspase 3
Cdk: cyclin dependent kinase
CK: cytokeratin
ERα: estrogen receptor alpha
MaSC: mammary stem cell
MEC: mammary epithelial cell
PR: progesterone receptor
SFE: sphere forming efficiency
SMA: smooth muscle actin

## Declarations

### Ethics Approval and Consent to Participate

Mice were maintained following the Canadian Council of Animal Care guidelines under animal utilization protocol 20-15 approved by the University of Windsor.

### Consent for Publication

Not applicable

### Data Availability Statement

Data is available upon request.

### Author Contribution

BF, ERA and LAP contributed to the project design. BF, JV and EH contributed to data acquisition. BF, JV, EH, ERA and LAP contributed to the data analysis. BF and LAP prepared the manuscript. LAP secured the funding for this study. All authors read and approved the final manuscript.

## Acknowledgements

We thank Drs. C. Pin, F. Dick and L. Drysdale from the London Regional Transgenic and Gene Targeting Facility for the transgenic injections and helpful advice. Special thanks to Dr. W. Muller for the MMTV-SV40-TRPS-1 vector, the University of Windsor Central Animal Care Facility and University of Windsor Flow Cytometry Facility. Thanks to E. Fidalgo da Silva, D. Lubanska, I. Hinch and N. Philbin for assistance with idea generation, genotyping and technical assistance.

