## Supplemental Files for "Atypical Cell Cycle Regulation Promotes Mammary Stem Cell Expansion and Therapeutic Resistance"

### Figure S1

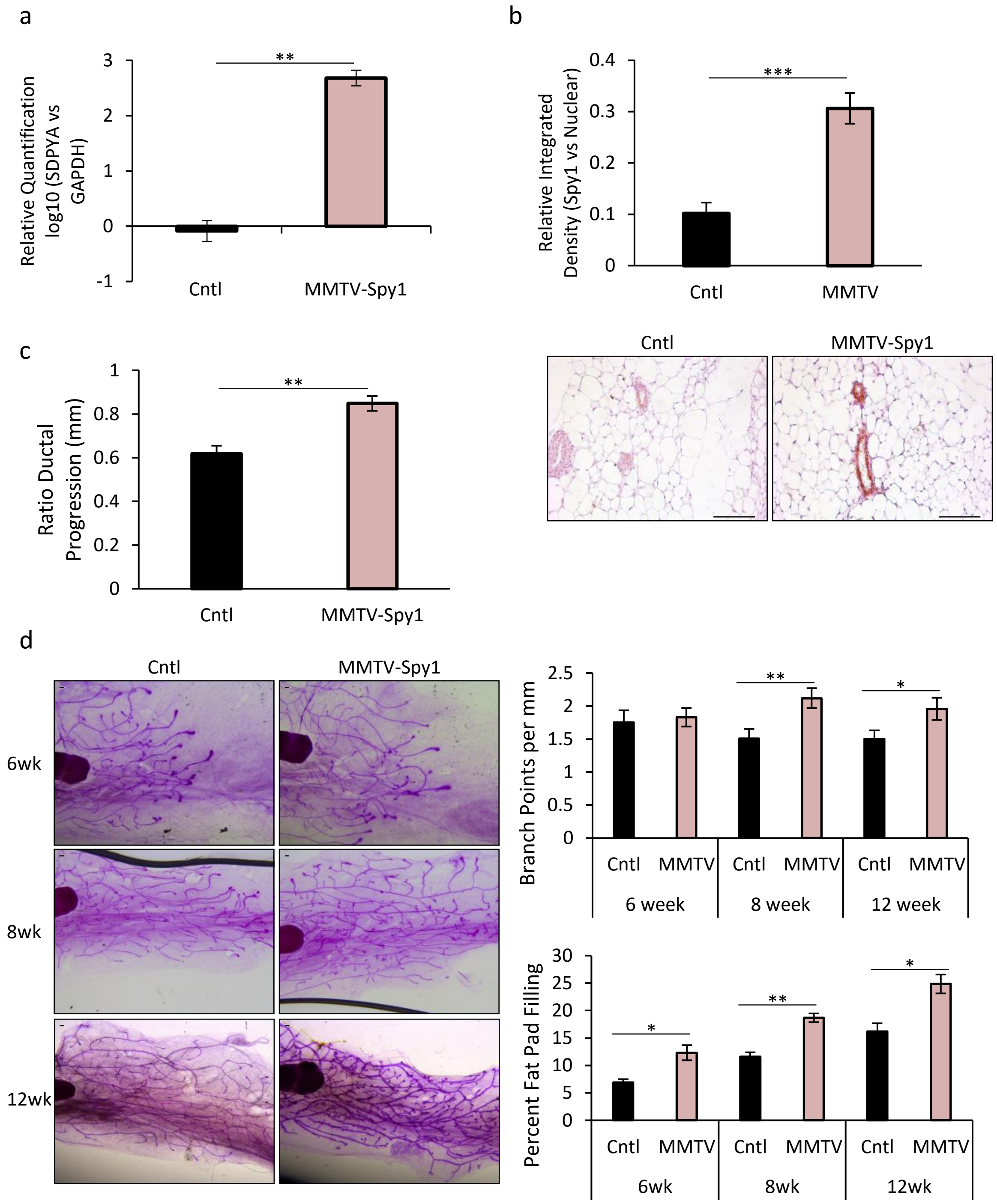

a
Relative Quantification log10 (SDPYA vs GAPDH)
3
2
1
0
-1
**
Cntl
MMTV-Spy1
b
Relative Integrated Density (Spy1 vs Nuclear)
0.4
0.3
0.2
0.1
0
***
Cntl
MMTV
c
Ratio Ductal Progression (mm)
1
0.8
0.6
0.4
0.2
0
**
Cntl
MMTV-Spy1
Cntl
MMTV-Spy1
d
Cntl
MMTV-Spy1
6wk
8wk
12wk
Branch Points per mm
2.5
2
1.5
1
0.5
0
**
*
Cntl
MMTV
6 week
8 week
12 week
Percent Fat Pad Filling
30
25
20
15
10
5
0
*
**
*
Cntl
MMTV
6wk
8wk
12wk

Figure S2

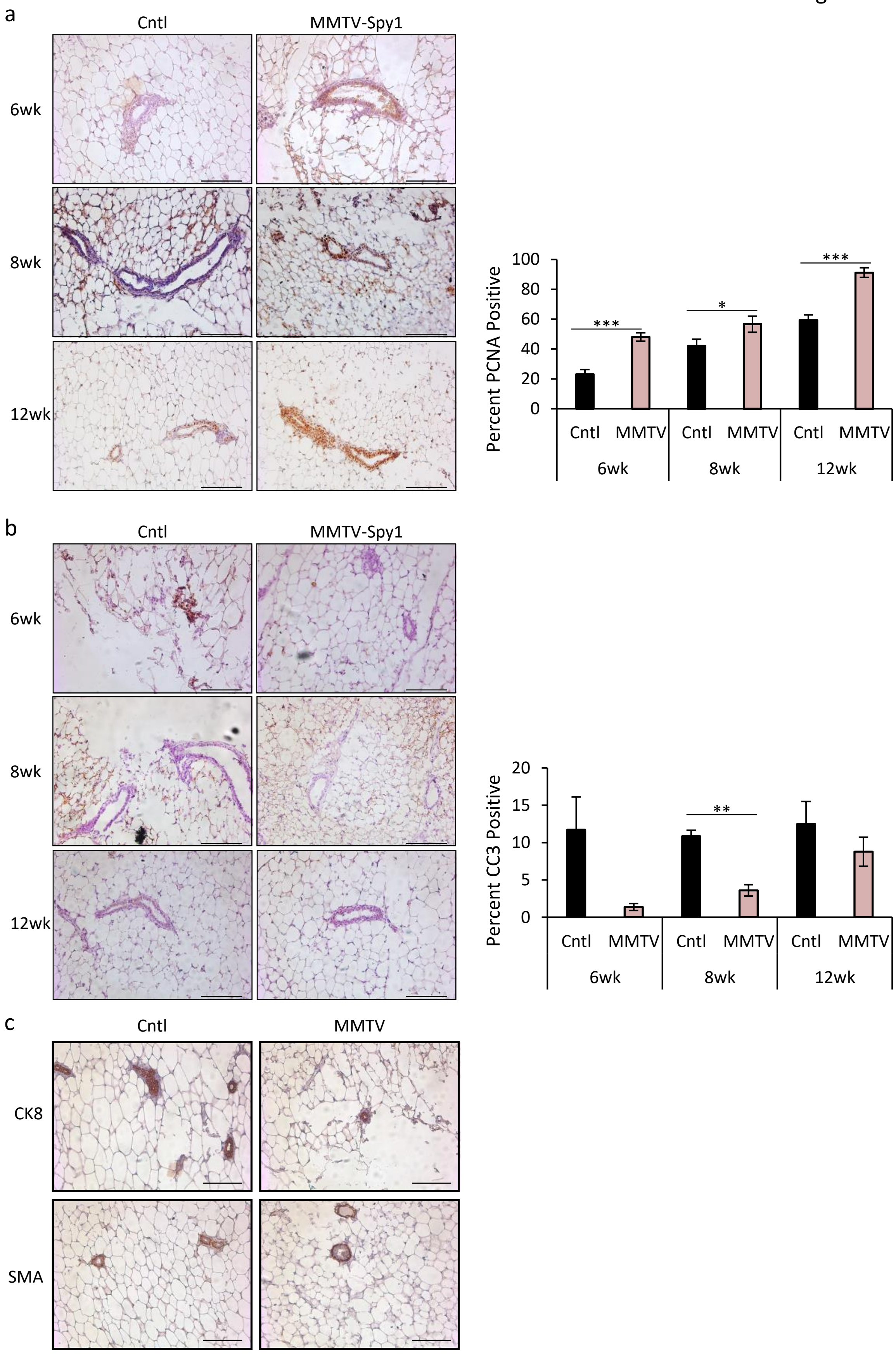

a
Cntl
MMTV-Spy1
6wk
8wk
12wk
Percent PCNA Positive
100
80
60
40
20
0
***
*
***
Cntl
MMTV
Cntl
MMTV
Cntl
MMTV
6wk
8wk
12wk
b
Cntl
MMTV-Spy1
6wk
8wk
12wk
Percent CC3 Positive
20
15
10
5
0
**
Cntl
MMTV
Cntl
MMTV
Cntl
MMTV
6wk
8wk
12wk
c
Cntl
MMTV
CK8
SMA

Figure S3

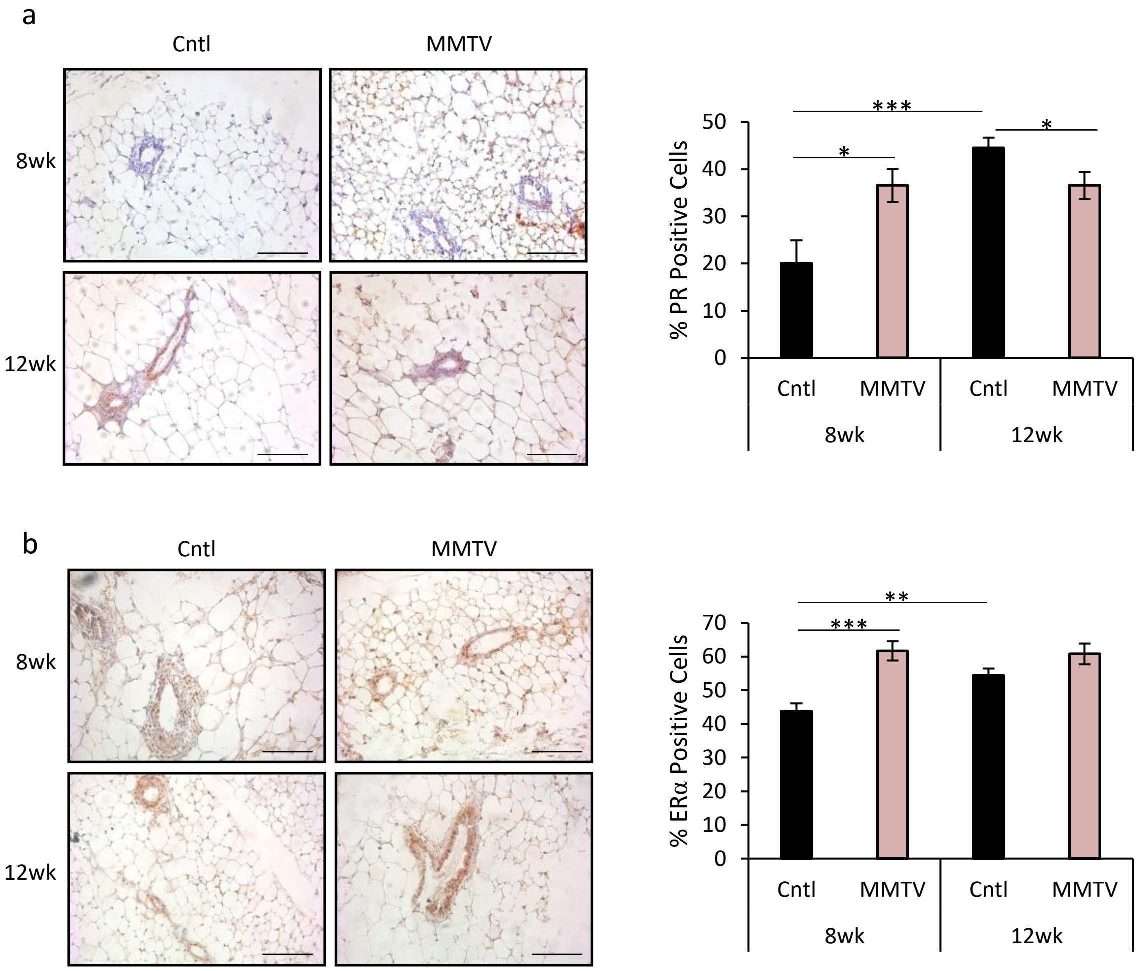

a
Cntl
MMTV
8wk
12wk
% PR Positive Cells
50
40
30
20
10
0
***
*
*
Cntl
MMTV
Cntl
MMTV
8wk
12wk
b
Cntl
MMTV
8wk
12wk
% ERα Positive Cells
70
60
50
40
30
20
10
0
**
***
Cntl
MMTV
Cntl
MMTV
8wk
12wk

Figure S4

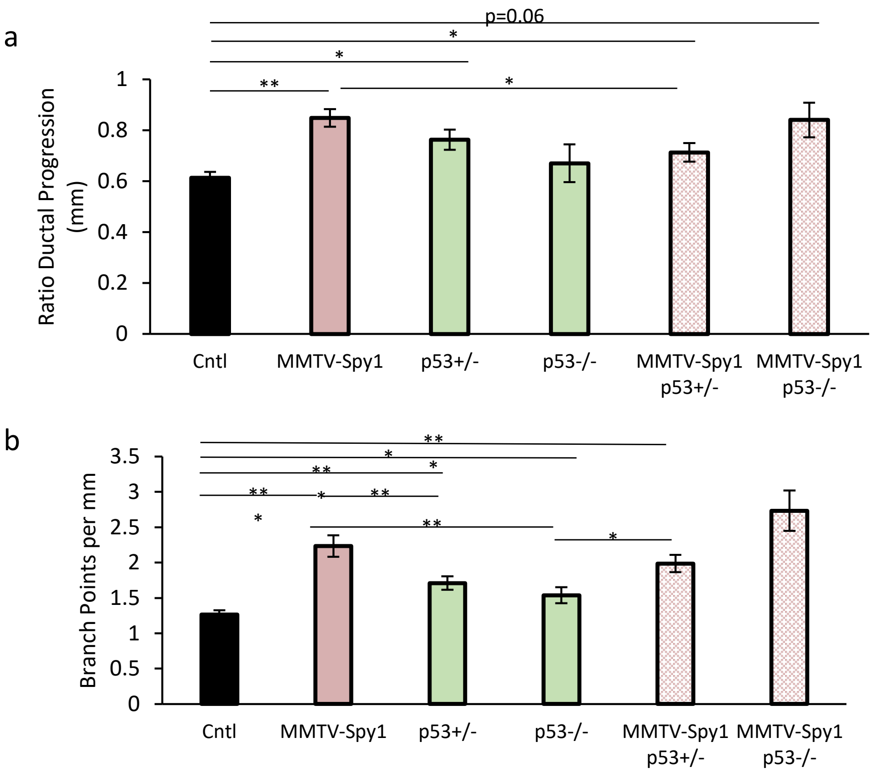
a
p=0.06
*
*
**
*
Ratio Ductal Progression (mm)
1
0.8
0.6
0.4
0.2
0
Cntl
MMTV-Spy1
p53+/-
p53-/-
MMTV-Spy1 p53+/-
MMTV-Spy1 p53-/-
b
**
*
**
*
**
*
**
*
**
*
Branch Points per mm
3.5
3
2.5
2
1.5
1
0.5
0
Cntl
MMTV-Spy1
p53+/-
p53-/-
MMTV-Spy1 p53+/-
MMTV-Spy1 p53-/-

Figure S5

a

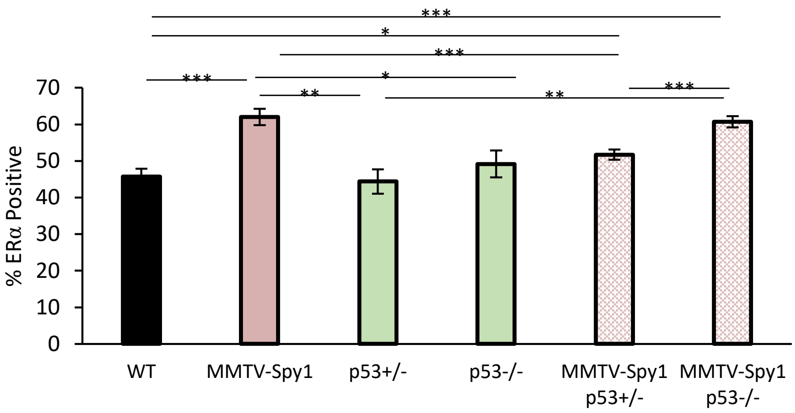

***
*
***
***
**
*
**
***
70
60
50
40
30
20
10
0
% ERα Positive
WT
MMTV-Spy1
p53+/-
p53-/-
MMTV-Spy1 p53+/-
MMTV-Spy1 p53-/-


b

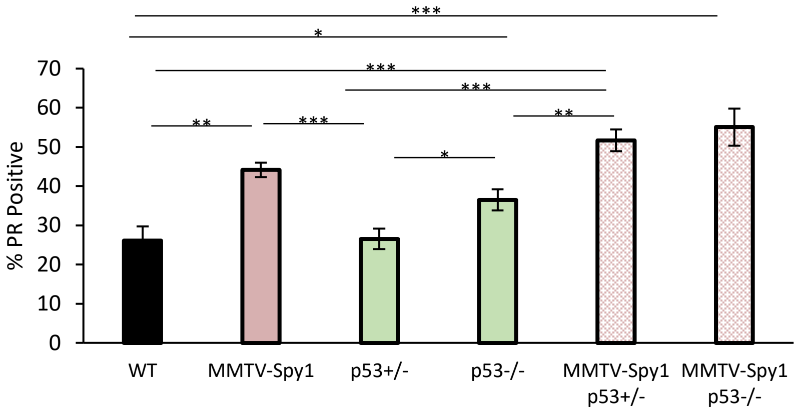

***
*
***
***
**
***
**
*
70
60
50
40
30
20
10
0
% PR Positive
WT
MMTV-Spy1
p53+/-
p53-/-
MMTV-Spy1 p53+/-
MMTV-Spy1 p53-/-

Figure S6

a

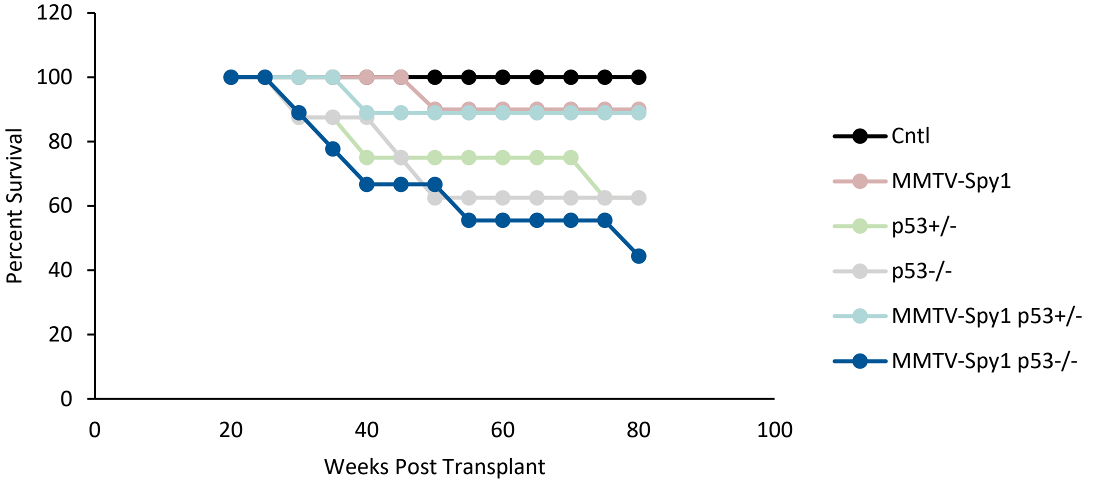
120
100
80
60
40
20
0
Percent Survival
0
20
40
60
80
100
Weeks Post Transplant
Cntl
MMTV-Spy1
p53+/-
p53-/-
MMTV-Spy1 p53+/-
MMTV-Spy1 p53-/-

b

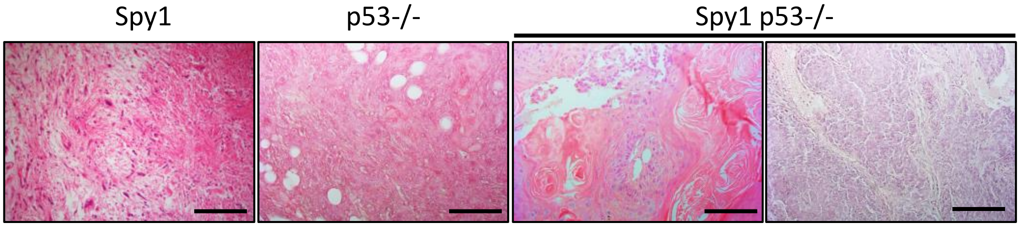
Spy1
p53-/-
Spy1 p53-/-

Figure S7

a

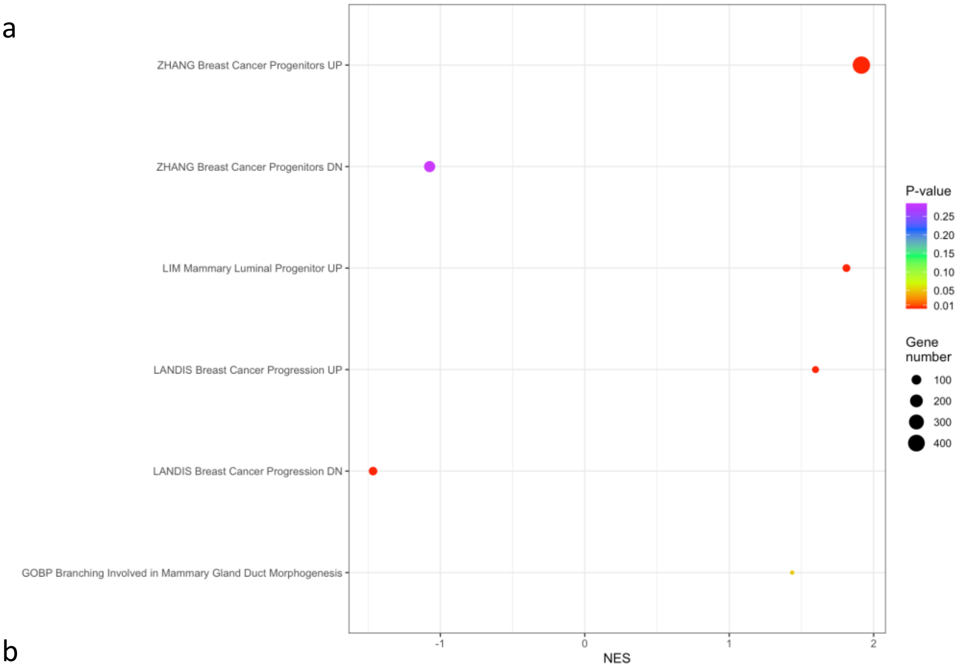
ZHANG Breast Cancer Progenitors UP
ZHANG Breast Cancer Progenitors DN
LIM Mammary Luminal Progenitor UP
LANDIS Breast Cancer Progression UP
LANDIS Breast Cancer Progression DN
GOBP Branching Involved in Mammary Gland Duct Morphogenesis
-1
0
1
2
NES
P-value
0.25
0.20
0.15
0.10
0.05
0.01
Gene number
100
200
300
400

b

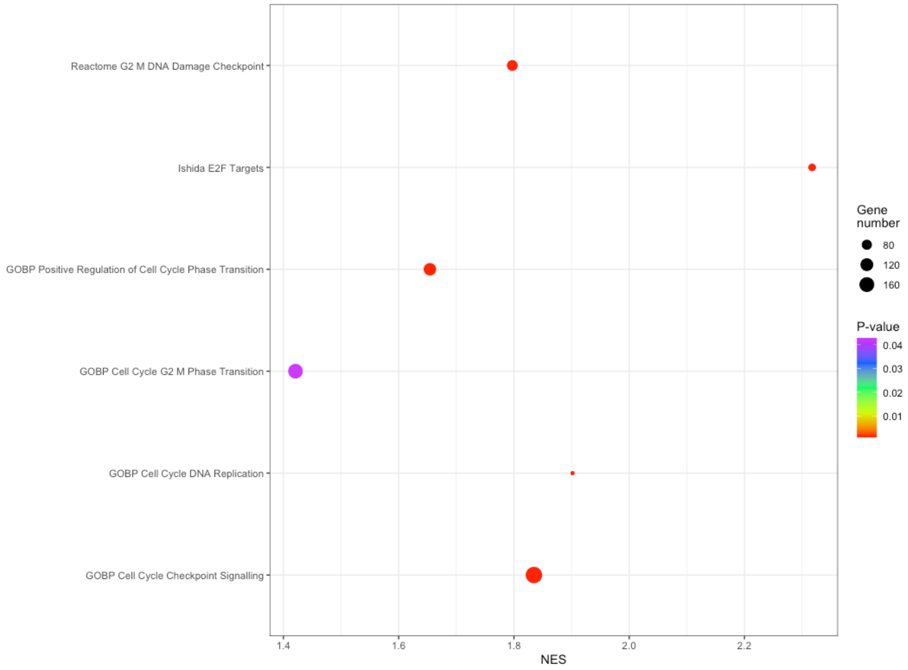
Reactome G2 M DNA Damage Checkpoint
Ishida E2F Targets
GOBP Positive Regulation of Cell Cycle Phase Transition
GOBP Cell Cycle G2 M Phase Transition
GOBP Cell Cycle DNA Replication
GOBP Cell Cycle Checkpoint Signalling
1.4
1.6
1.8
2.0
2.2
NES
Gene number
80
120
160
P-value
0.04
0.03
0.02
0.01

Figure S8

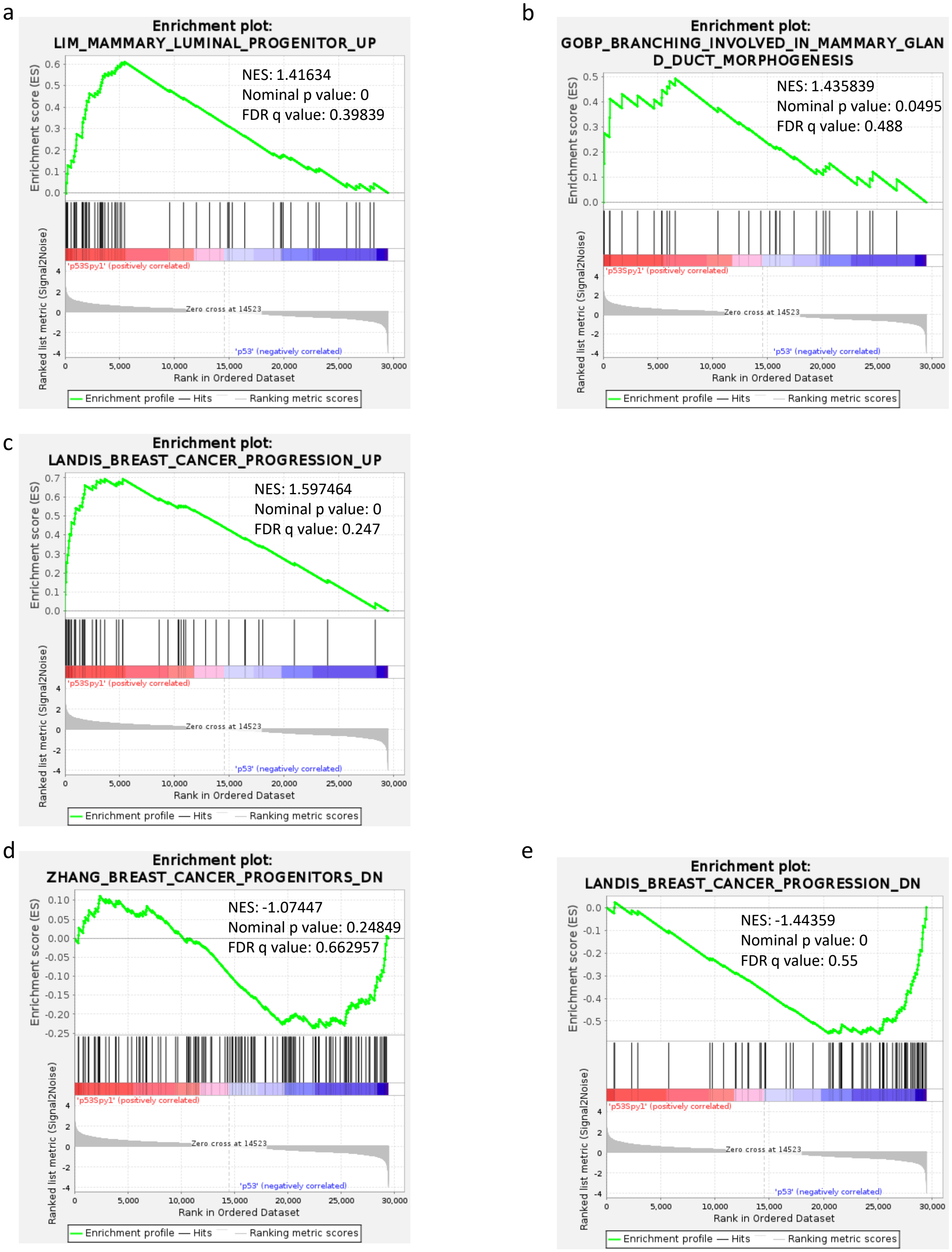
a
Enrichment plot: LIM_MAMMARY_LUMINAL_PROGENITOR_UP
NES: 1.41634
Nominal p value: 0
FDR q value: 0.39839
b
Enrichment plot: GOBP_BRANCHING_INVOLVED_IN_MAMMARY_GLAND_DUCT_MORPHOGENESIS
NES: 1.435839
Nominal p value: 0.0495
FDR q value: 0.488
c
Enrichment plot: LANDIS_BREAST_CANCER_PROGRESSION_UP
NES: 1.597464
Nominal p value: 0
FDR q value: 0.247
d
Enrichment plot: ZHANG_BREAST_CANCER_PROGENITORS_DN
NES: -1.07447
Nominal p value: 0.24849
FDR q value: 0.662957
e
Enrichment plot: LANDIS_BREAST_CANCER_PROGRESSION_DN
NES: -1.44359
Nominal p value: 0
FDR q value: 0.55
Enrichment score (ES)
Ranked list metric (Signal2Noise)
'p53Spy1' (positively correlated)
'p53' (negatively correlated)
Zero cross at 14523
Rank in Ordered Dataset
Enrichment profile
Hits
Ranking metric scores

Figure S9

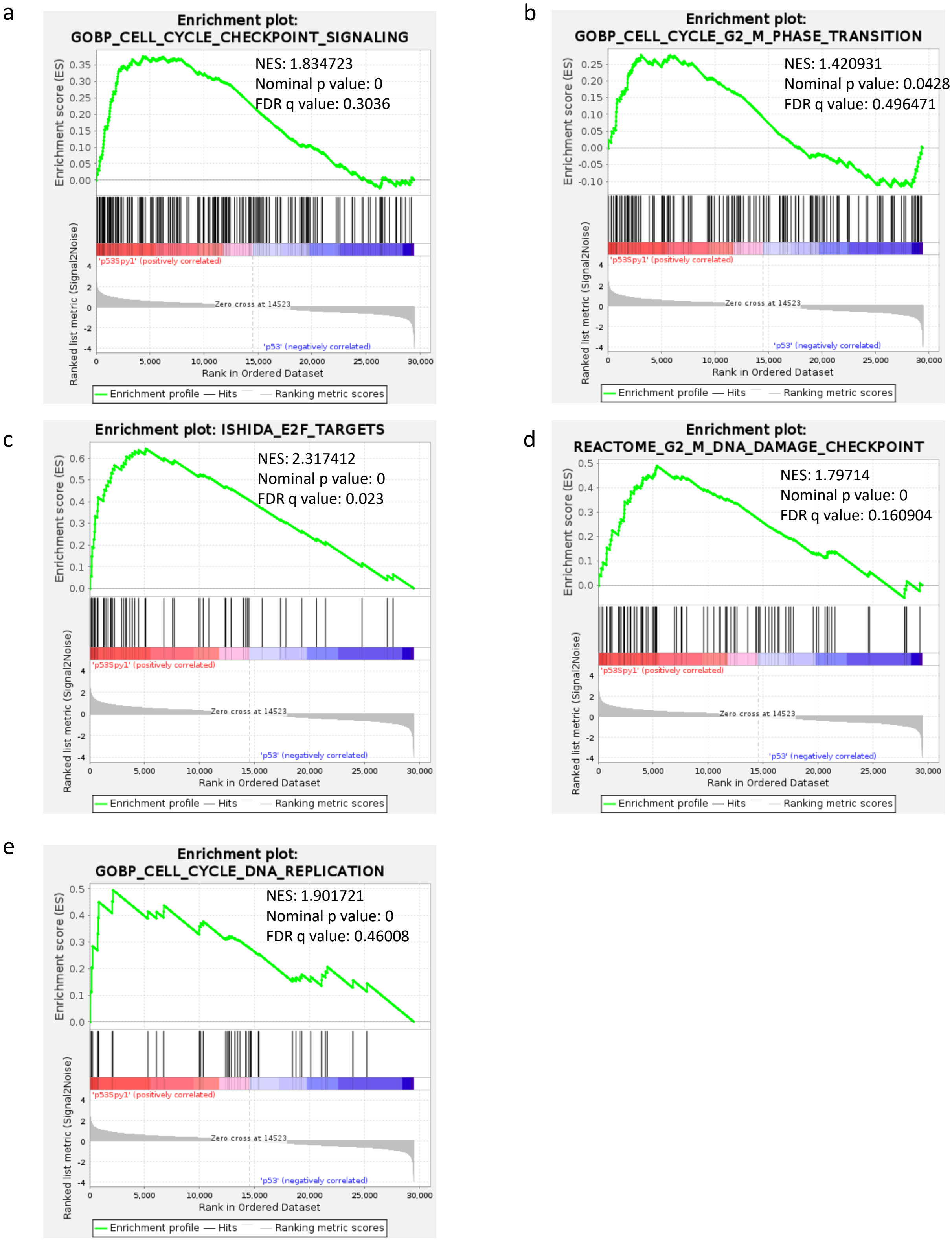
a
Enrichment plot: GOBP_CELL_CYCLE_CHECKPOINT_SIGNALING
NES: 1.834723
Nominal p value: 0
FDR q value: 0.3036
b
Enrichment plot: GOBP_CELL_CYCLE_G2_M_PHASE_TRANSITION
NES: 1.420931
Nominal p value: 0.0428
FDR q value: 0.496471
c
Enrichment plot: ISHIDA_E2F_TARGETS
NES: 2.317412
Nominal p value: 0
FDR q value: 0.023
d
Enrichment plot: REACTOME_G2_M_DNA_DAMAGE_CHECKPOINT
NES: 1.79714
Nominal p value: 0
FDR q value: 0.160904
e
Enrichment plot: GOBP_CELL_CYCLE_DNA_REPLICATION
NES: 1.901721
Nominal p value: 0
FDR q value: 0.46008
Enrichment score (ES)
Ranked list metric (Signal2Noise)
'p53Spy1' (positively correlated)
Zero cross at 14523
'p53' (negatively correlated)
Rank in Ordered Dataset
Enrichment profile — Hits — Ranking metric scores

Figure S10

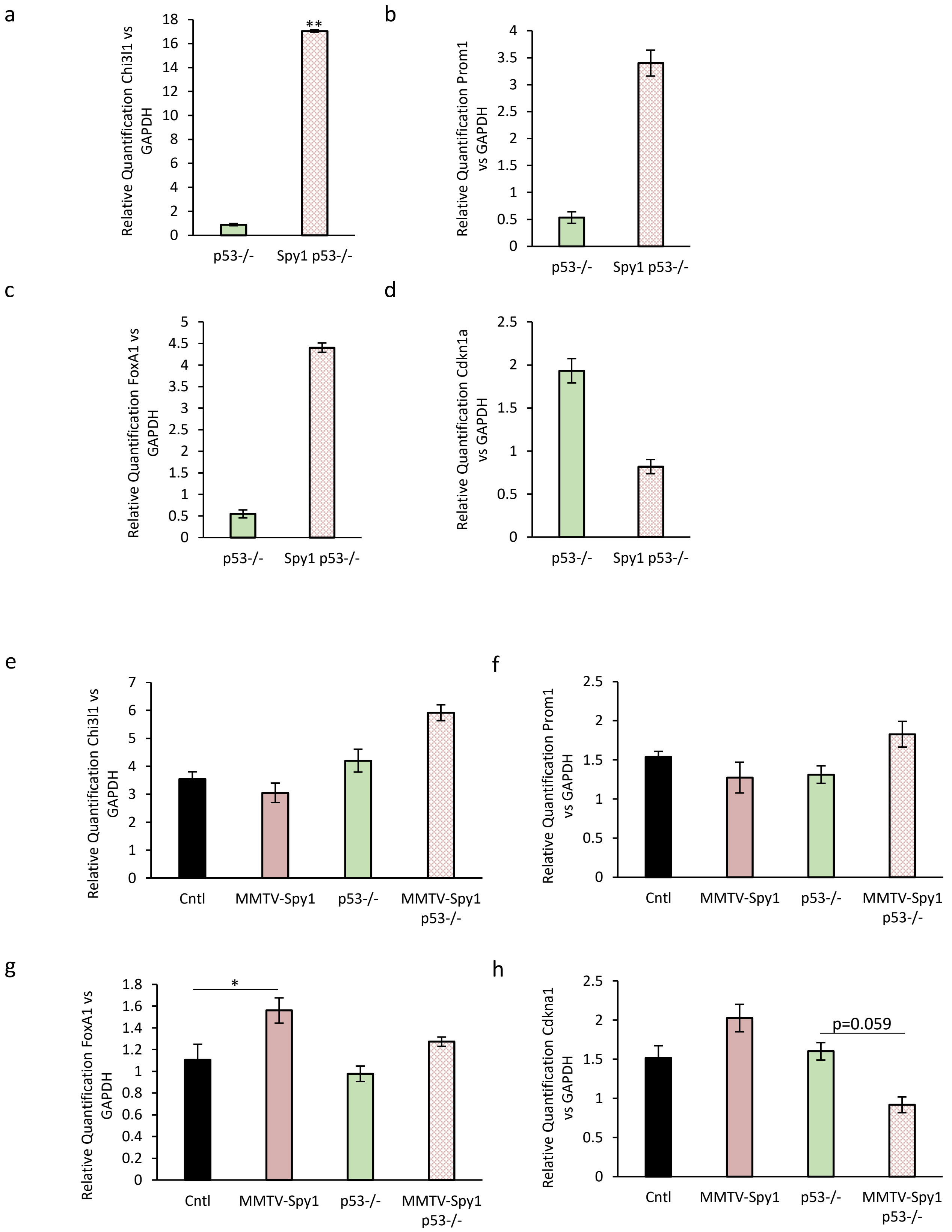
a
Relative Quantification Chi3l1 vs GAPDH
18
16
14
12
10
8
6
4
2
0
**
p53-/-
Spy1 p53-/-
b
Relative Quantification Prom1 vs GAPDH
4
3.5
3
2.5
2
1.5
1
0.5
0
p53-/-
Spy1 p53-/-
c
Relative Quantification FoxA1 vs GAPDH
5
4.5
4
3.5
3
2.5
2
1.5
1
0.5
0
p53-/-
Spy1 p53-/-
d
Relative Quantification Cdkn1a vs GAPDH
2.5
2
1.5
1
0.5
0
p53-/-
Spy1 p53-/-
e
Relative Quantification Chi3l1 vs GAPDH
7
6
5
4
3
2
1
0
Cntl
MMTV-Spy1
p53-/-
MMTV-Spy1 p53-/-
f
Relative Quantification Prom1 vs GAPDH
2.5
2
1.5
1
0.5
0
Cntl
MMTV-Spy1
p53-/-
MMTV-Spy1 p53-/-
g
Relative Quantification FoxA1 vs GAPDH
1.8
1.6
1.4
1.2
1
0.8
0.6
0.4
0.2
0
*
Cntl
MMTV-Spy1
p53-/-
MMTV-Spy1 p53-/-
h
Relative Quantification Cdkna1 vs GAPDH
2.5
2
1.5
1
0.5
0
p=0.059
Cntl
MMTV-Spy1
p53-/-
MMTV-Spy1 p53-/-

Figure S11

a

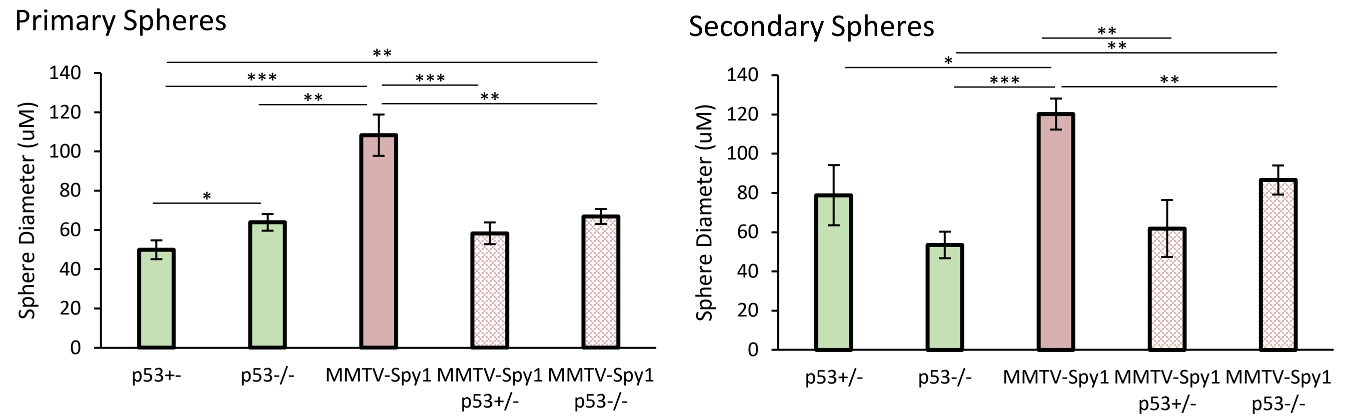
Primary Spheres
Sphere Diameter (uM)
140
120
100
80
60
40
20
0
*
**
***
**
***
**
p53+-
p53-/-
MMTV-Spy1
MMTV-Spy1 p53+/-
MMTV-Spy1 p53-/-
Secondary Spheres
Sphere Diameter (uM)
140
120
100
80
60
40
20
0
**
**
*
***
**
p53+/-
p53-/-
MMTV-Spy1
MMTV-Spy1 p53+/-
MMTV-Spy1 p53-/-

#### Supplemental Figure Legends

**Figure S1**: Spy1 increases ductal branching. **a**) qRT-PCR analysis of inguinal glands from MMTV-Spy1 and littermate controls (Cntl) for Spy1 levels corrected for GAPDH (n=4). **b**) Spy1 expression in inguinal mammary glands collected from 8-week-old MMTV-Spy1 and littermate control (Cntl) mice. Representative images in bottom panels with quantification of Spy1 levels using ImageJ software analysis shown in the top panel. Scale bar = 100 μm (n=3). **c**) Ratio of ductal progression past the lymph node in 8-week-old inguinal mammary glands from control littermates (Cntl) and MMTV-Spy1 mice (n=3). **d**) Whole mount analysis of inguinal mammary glands from 6-, 8- and 12-week-old littermate control and MMTV-Spy1 mice. Representative images shown in left panels, and quantification of number of branch points per mm (top right panel) and percentage of fat pad filling (bottom right panel) were quantified using ImageJ software analysis. Scale bar= 0.1mm. (6-week n=4-, 8- and 12-week n=3); Errors bars represent SE; Student's T-test. *p<0.05, **p<0.01, ***p<0.001.

**Figure S2:** Spy1 increases proliferation and decreases apoptosis. **a**) PCNA expression in MMTV-Spy1 and littermate controls via immunohistochemical analysis. Quantification of percentage of PCNA positive mammary epithelial cells over five fields of view per sample. Scale bar=100μM **b**) Cleaved caspase-3 (CC3) expression in MMTV-Spy1 and littermate controls via immunohistochemical analysis. Quantification of percentage of CC3-positive mammary epithelial cells over five fields of view per sample. Scale bar=100μM. **c**) Representative images of immunohistochemical analysis of CK8 and SMA. Scale bar=100μM. n=3; Error bars represent SE; Student's T test. *p < 0.05, **p < 0.01, ***p < 0.001.

**Figure S3:** Altered hormone receptor status with elevated Spy1. Immunohistochemical analysis of **a**) PR and **b**) ER in 8- and 12-week-old MMTV-Spy1 and littermate control inguinal mammary glands. Representative images shown in left panels, where blue represents hematoxylin nuclear stain and brown represents **a**) PR or **b**) ER. Quantification of the percent positive **a**) PR and **b**) ER cells are depicted in right panels. Scale bar = 100 µm; n=3; Errors bars represent SE; Student's T-test. *p<0.05, **p<0.01, ***p<0.001.

**Figure S4:** Loss of p53 does not alter Spy1 driven mammary development effects. Whole mount analysis was performed on inguinal mammary glands from 8 week old MMTV-Spy1 and p53 null intercrossed mice. **a**) Ductal progression past lymph node and **b**) number of branch points per mm of primary duct was quantified (Cntl n=4, MMTV-Spy1 n=5, p53+/- n=5, p53-/- n=4, MMTV-Spy1 p53+/- n=7, MMTV-Spy1 p53-/- n=3). Scale bar=100µM. n=3; Error bars represent SE; Student's T test. *p < 0.05, **p < 0.01, ***p < 0.001.

**Figure S5:** Loss of p53 does not alter hormone receptor expression in MMTV-Spy1 mice. Immunohistochemical analysis of the percent positive **a**) ER and **b**) PR in 8-week-old inguinal mammary glands of MMTV-Spy1 and p53 null intercrossed mice. Percent positive cells is represented graphically (Cntl n=6, MMTV-Spy1 n=7, p53+/- n=4, p53-/- n=3, MMTV-Spy1 p53+/- n=6, MMTV-Spy1 p53-/- n=3). Errors bars represent SE; Student's T-test. *p<0.05, **p<0.01, ***p<0.001.

**Figure S6:** Primary mammary epithelial cells from MMTV-Spy1 and p53 null intercrossed mice were injected into the cleared fat pads of wildtype mice and monitored weekly for HANs and tumour development. **a**) Timing of tumour onset is depicted. **b**) Representative hematoxylin and eosin stained images from Spy1, p53 null and Spy1 p53 null tumours. Scale bar=100µM. (Cntl n=15, MMTV-Spy1 n=17, p53+/- n=21, p53-/- n=14, MMTV-Spy1 p53+/- n=21, MMTV-Spy1 p53-/- n=13).

**Figure S7**: Dot plots depicting 2 sets of gene sets altered in Spy1 p53 null tumours as compared to p53 null tumours. Plots depict altered **a**) breast specific gene sets and **b**) cell cycle progression gene sets.

**Figure S8**: Spy1 driven tumours have enrichment in progenitor and breast cancer progression signaling. GSEA demonstrated that Spy1 p53 null tumours are enriched in **a**) mammary luminal progenitors, **b**) branching morphogenesis, and **c**) breast cancer progression gene sets and are depleted for genes downregulated in **d**) breast cancer progenitors and **e**) breast cancer progression.

**Figure S9**: Spy1 driven tumours are enriched in cell cycle progression gene sets. GSEA demonstrated that Spy1 p53 null driven tumours are enriched in **a**) cell cycle checkpoint signaling, **b**) cell cycle G2 M phase transition, **c**) E2F targets, **d**) G2 M DNA damage checkpoint, and **e**) cell cycle DNA replication gene sets.

**Figure S10**: Changes in expression in Spy1 driven tumours are reflected during normal development. qRT-PCR analysis of expression of **a**) Chi3l1, **b**) Prom1, **c**) FoxA1 and **d**) Cdkn1a corrected for GAPDH in Spy1 p53 null and p53 null tumours (p53 null n=3; Spy1 p53 null n=5). qRT-PCR analysis of expression of **e**) Chil3l1, **f**) Prom1, **g**) FoxA1 and **h**) Cdkn1a corrected for GAPDH from inguinal mammary glands of 8 week old MMTV-Spy1 p53 null intercrossed mice (n=3). Errors bars represent SE; Student's T-test. *$p<0.05$, **$p<0.01$.

**Figure S11: a**) Primary tumour cells from MMTV-Spy1 and p53 null tumours were cultured as mammospheres. Sphere diameter of primary (left panel) and secondary (right panel) mammospheres was quantified. N=3; Error bars represent SE; Student's T test. *$p < 0.05$, **$p < 0.01$, ***$p < 0.001$.
